# Measurable fields-to-spike causality and its dependence on cortical layer and area

**DOI:** 10.1101/2023.01.17.524451

**Authors:** Shailaja Akella, André M. Bastos, Earl K. Miller, Jose C. Principe

## Abstract

Distinct dynamics in different cortical layers are apparent in neuronal and local field potential (LFP) patterns, yet their associations in the context of laminar processing have been sparingly analyzed. Here, we study the laminar organization of spike-field causal flow within and across visual (V4) and frontal areas (PFC) of monkeys performing a visual task. Using an event-based quantification of LFPs and a directed information estimator, we found area and frequency specificity in the laminar organization of spike-field causal connectivity. Gamma bursts (40-80 Hz) in the superficial layers of V4 largely drove intralaminar spiking. These gamma influences also fed forward up the cortical hierarchy to modulate laminar spiking in PFC. In PFC, the direction of intralaminar information flow was from spikes → fields where these influences dually controlled top-down and bottom-up processing. Our results, enabled by innovative methodologies, emphasize the complexities of spike-field causal interactions amongst multiple brain areas and behavior.

## Introduction

Whereas correlation models describe the dependence structure between observed variables, causal models go one step further: they predict whether a variable directly affects a change in another. Understanding the causal interactions between neural entities is central to studying brain function. In this pursuit, several statistical methods have been developed at each scale of the electrophysiological measures - spiking activity and local field potentials (LFPs). In LFPs, Granger causality studies have implicated distinct frequency bands in several neurophysiological phenomena, including predictive routing (*1, 2*), top-down and bottom-up signal processing (*3–6*), and working memory (*7, 8*). Pairwise correlations between single neurons have identified network organization consistent with anatomical connectivity (*9–13*), detected distinct functionally specialized populations (*13, 14*), and enabled foundational understanding of behavior encoding processes (*15–17*).

However, studies in individual signal modalities only partially explain brain function. A single neuron cannot support complex behavior like perception, cognition, and action. These capabilities emerge only when networks of neural circuits, as a whole, define specific brain functions. Because LFPs account for the joint activity of neuronal assemblies, they constitute a critical ‘middle ground’ between single neuron activity and behavior (*18, 19*). To that end, bridging this pathway from brain structure to function requires detailed knowledge of the effective influences across the signal modalities. Yet, the distinct class of spike-field causal analyses is not without challenges. Firstly, spike trains are discrete binary-valued signals modeled as point processes, whereas field potentials are continuous signals preferably analyzed using classical signal processing tools. Secondly, compared with the millisecond dynamics of spiking activity, LFP dynamics vary much too slowly at orders of a few hundred milliseconds. Lastly, it is difficult to completely dissociate spiking-related events from LFPs, eventually contributing to spurious associations between the signals (*20*). These inequalities in the multiscale brain activity call for specialized methodologies that must explicitly incorporate the concepts of neurophysiological signal generation, time resolution in the tens of milliseconds, and a shared metric space between the modalities.

Any construct of an effective cortical network must consider the influence of the anatomical parcellations of its structure. Anatomical tracing studies in the cortex have found that the laminar location of cell bodies and their termination patterns underlies a hierarchical organization of cellular connections (*21, 22*). In other words, its position in the hierarchy affects the diversity of synaptic inputs a neuron receives, influencing its functionality. For instance, in many parts of the cortex, areas higher up the hierarchy represent increasingly complex features of the stimuli (visual, (*23–25*); auditory, (*26, 27*); somatosensory, (*28*)). Even globally, feedforward and feedback connections differentially recruit neuronal populations in a layer-specific manner (*2, 29–31*). Likewise, quantitative modeling efforts of such a hierarchical flow have generated testable predictions. Most noted is the microcircuitry developed by Douglas and Martin (*32*). Based on their studies in the cat-visual cortex, they presented a conduction-based connectivity model between the thalamic and cortical neurons. These results have since formed the basis for canonicity over the cortical sheet. Another prediction borne out of hierarchical quantitative modeling is that the brain may employ empirical Bayesian inference on its sensory inputs (*33–35*), providing support for several neural computation theories, including predictive coding, hierarchical message passing, and mean-field interactions.

Although recent technological advancements in electrode design have enabled simultaneous measurement of neurons and LFPs at multiple, interconnected hierarchical levels, only a few studies have explored their associations while accounting for the hierarchical cortical processes. Noteworthy analyses include those conducted by (*36*). Here, during a task of directed spatial attention, primate visual areas 1, 2, and 4 (V1, V2, and V4, respectively) reported significantly different spike-field coherence patterns between the superficial and deep layers. Single neurons coherent with gamma band (40 − 60 Hz) frequencies were confined to the superficial layers, while those synchronized with alpha band frequencies (6−16 Hz) were dominantly found in the deep layers. In areas V2 and V4, spike-field coherence was modulated by attention - decreases in alpha synchrony contrasted with increases in gamma synchrony. While these results set precedence for spike-field association studies in the cortical column, the implemented metrics allowed for correlative but not causal inferences. Further, spectral estimations of LFPs limit access to timing information important to deciphering immediate causal influences between the signals. Lastly, their findings only focused on the visual cortex.

In this work, we address the above challenges by implementing a neurophysiologically relevant interpretation of the LFP traces in terms of their oscillatory burst patterns. Characterized by their timing, amplitude, and duration, the bursts collectively summarize a marked point process representation of the LFP traces (*37*) - therefore, bringing the two modalities into the same algebraic space of point processes while maintaining the fine temporal structure of the oscillations. Next, we introduce an information theoretic metric, directed information, in the point process space enabling causal inferences between the signal modalities. To analyze the specifics of feedforward and feedback spike-field connectivity, we apply our methodologies on layer-resolved, simultaneous recordings of LFPs and spiking activity from the visual and frontal areas of two primates performing a visual task with modulated stimulus predictability. We build on previous analyses conducted on the same dataset (*2*) where causal connectivity between LFPs revealed interesting asymmetric networks in the beta and gamma frequency ranges. Accordingly, we focus our analyses of spike-field causal connectivity with gamma bursts in the 40 − 80 Hz frequency range and beta bursting in a broad frequency range of 8 − 30 Hz. These analyses revealed layer and frequency-specific asymmetries in spike-field associations and presented causal evidence for the direction of flow of these signals. The findings suggest a complex dynamic between the two modalities consistent in frequency and layer asymmetries while revealing a complicated directional influence pattern.

## Results

### Projection of field potentials onto the point process space

Recognizing the need for simultaneous studies of spiking activity and local field potentials (LFPs), we propose to analyze causal connectivity patterns between the multiscale activity using an event-based quantification of LFPs. Specifically, we summarize each LFP trace as a point process of transient bursts of narrowband oscillations in established frequency ranges (beta, 8 - 30 Hz, and gamma, 40 - 80 Hz). The resulting representation allows for a unified metric space between spiking activity and LFPs while preserving the time resolution across the different recording modalities. To realize such a representation, we developed an unsupervised learning model named the ‘generative model for oscillatory bursts’ (*37*). Unlike other time-frequency decomposition techniques, the generative model explicitly incorporates the concepts of neurophysiological signal generation and enables a high-time-resolution representation of transient activity. The model achieves this by isolating oscillatory waveforms from the spontaneous background of bandpass LFPs using sparse coding techniques. First, we divide the bandpass LFP data into training and test sets. On the training data, identified oscillatory waveforms are compiled into a dictionary of representative waveforms. The final point process representation of the test signals is then obtained by a convolution between each bandpass LFP trace and the learned dictionary using adaptive thresholding on the burst power (Figure 1**A**).

**Figure 1:**
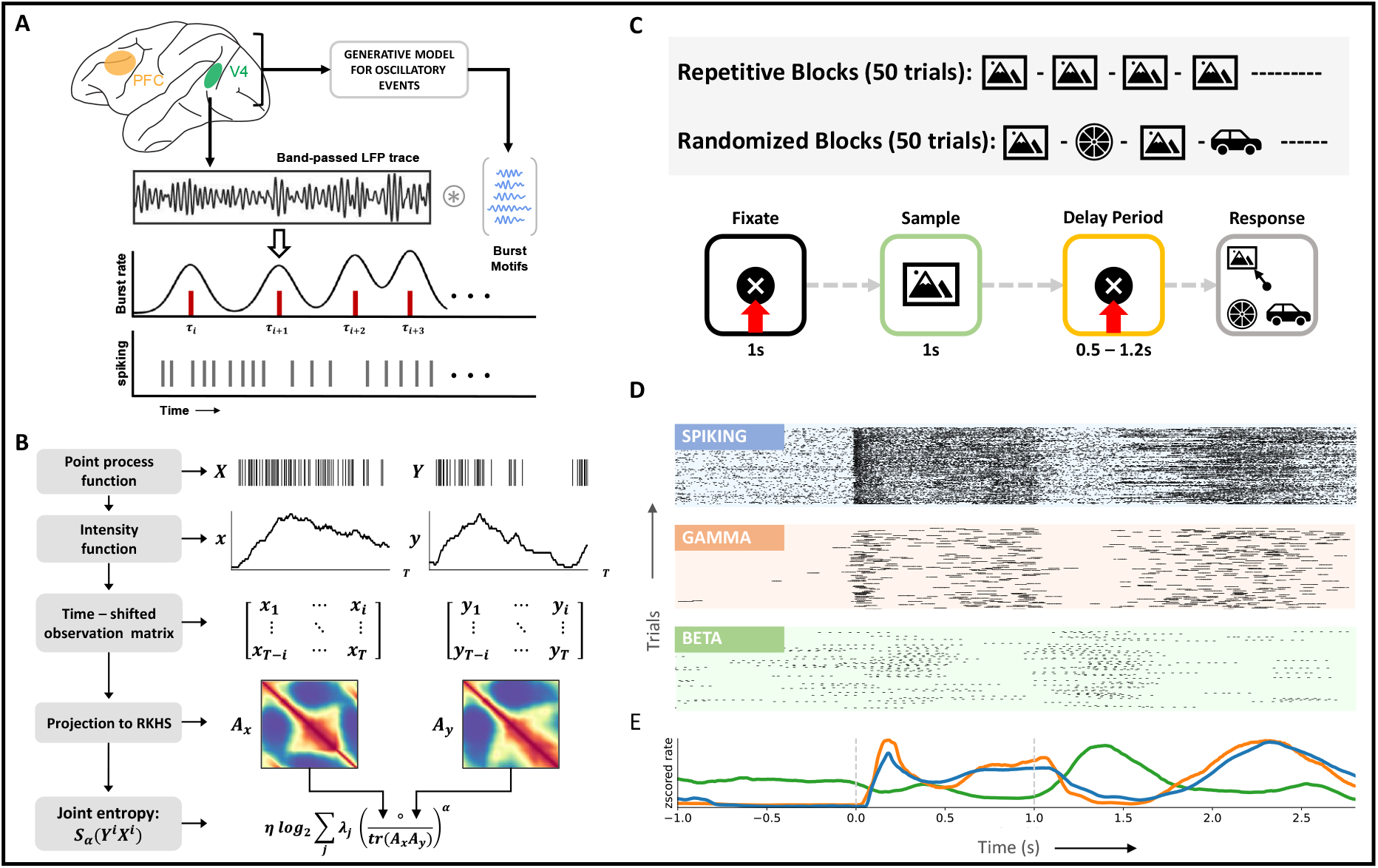
Methods and behavioral task. **A,** Mapping of local field potentials from continuous space to point process space. Using the generative model, a dictionary of typical oscillatory patterns is learnt from bandpassed LFP traces (training). Construction of the point process space for LFPs follows from a convolution between LFP traces and the learnt dictionary (testing).**B**, Example evaluation of joint entropy term in directed information (equation 16) using matrix formulation of Renyi’s entropy, *α*-entropies. At time *i*, each time-shifted spiking matrix is projected onto a high–dimensional RKHS defined by the kernel, *κ*. Eigen-spectrum of the Hadamard product between the normalized Gram matrices captures the joint entropy between the two time series. **C**, (Top) Experimental setup. Following a 1s-fixation period, the subject is shown a sample stimulus for 1s. At the end of a delay period (fixed/variable), the subject saccades to the sampled stimulus that reappears at one of four randomized locations along with distractor images. **D**, Exemplar spiking activity and LFPs from layer 2/3 of V4 visualized in the same space of point processes, subject 2. For visualization purposes, every peak of the detected oscillatory burst has been demarcated. **E**, Multi-scale average rate plots obtained via kernel smoothing methods, subject 2. All activity is aligned with the trial intervals in **C**.

To examine the role played by oscillatory dynamics in sensory information processing, we apply the generative model to laminar LFPs recorded from visual area 4 (V4) and prefrontal cortex (PFC) of two monkeys. During the recording, the monkeys actively engaged in a delayed match to sample (DMS) task (Figure 1**C**, bottom). In each session of the DMS task, the predictability of the stimulus was additionally modified every 50 trials in one of two ways: 1) the same stimulus was cued in consecutive trials during the ‘repetitive’ trial blocks, and 2) a randomly sampled stimulus was cued in consecutive trials during ‘randomized’ trial blocks (Figure 1**C**, top). Across ten sessions and two monkeys, we analyzed LFPs in the gamma (40 − 80 Hz) and beta (8 − 30 Hz) frequency ranges. Learning on the model was performed per session, area, and frequency range, culminating in 4 dictionaries per session (two areas and two frequency ranges). Example waveforms from each dictionary are summarized in Figure S1.

The main advantage of a point process representation of LFPs resides in its interpretability via rasters and rate plots. While the former supplies a platform for identifying precise timing-based modulations, the latter enables inferences based on the general properties of the point process. Figures 1**D, E** present representative rasters and rate functions over the complete trial duration for all signal modalities in area V4 (PFC, Figure S5). For better visualization, we plot rasters of beta and gamma bursts to demarcate each peak in an oscillatory burst. These plots succinctly summarize associations observed in previous studies (*2,38,39*) between spiking activity and gamma bursts (*r*_*V* 4_ = 0.66, *r*_*PF C*_ = 0.93, *p <* 0.05) and their anti-correlation with beta bursting (neuron, *r*_*V* 4_ = −0.62, *r*_*PF C*_ = −0.13, *p <* 0.05; gamma, *r*_*V* 4_ = −0.50, *r*_*PF C*_ = −0.19, *p* < 0.05). This anti-correlation between the low and high-frequency signals can be associated with the task. Increased spiking activity and gamma bursting are observed during bottom-up processing tasks, i.e., during sample and response intervals. Whereas activity in the beta frequency range is higher in the delay period, during which top-down processing streams are active.

To further validate the burst detections made by the generative model (Simulations, see Methods, Figure S3), we replicate results from a previous study (*2*) on the same dataset. For this, during the fixation and sample intervals, we analyzed the effect of stimulus predictability on gamma and beta power in V4 and PFC (Figure S2). Power fluctuations in each frequency range are determined by including the local burst power in the LFP point process’s rate function (see Methods, Burst power). The high time resolution of the LFP point process allows for sample-by-sample significance analysis of the signal local power enabling additional inference on the timing of power modulations. Throughout the sample interval, we observed increased gamma activity in V4 to ‘randomized’ stimuli (Figure S2**A**, right). Specifically, superficial layer gamma power in the early sample period noted more significant increases than the deep layer gamma power (p *<* 0.05, ANOVA). Gamma band power in the PFC was not modulated by sample predictability. Opposite effects were observed in the beta band power. In V4 and PFC, beta power increased during repeated stimulus presentations (p *<* 0.01, ANOVA). In this case, beta power modulations to sample predictability were more prominent in the deep layers than in the superficial layers across both areas (p < 0.05, ANOVA). This layer specialization of beta activity was observed only in the later half of the sample interval. In the fixation interval, the effect of stimulus predictability was most pronounced in the early periods, where beta power modulations in the deep layers of V4 were significantly higher (ANOVA, p < 0.01) during repeated presentations of the stimulus.

These results, combined with the results from (*2*), provide strong evidence for the involvement of oscillatory LFP rhythms in bottom-up and top-down information processing, where the separate pathways involving distinct cortical layers, timings, and frequencies point to a hierarchical system that mediates information integration.

### Visual stimulation reorganizes baseline network connectivity

Prior to studying the interactions between spiking activity and LFPs, we were interested in analyzing the interactivity at each scale. While previous work ((*2*)) has explored network interactions among LFPs, here, we study the dynamical changes in the V4 neuronal networks during sensory stimulation. To study these changes, we compared the interaction patterns between single neurons across the fixation and sample intervals. Specifically, we estimated directed information (DI) to identify causal influences between neurons in individual cortical columns of V4. DI is an information-theoretic measure that quantifies the magnitude and direction of information flow from one random variable to another. An improvement over mutual information (equation 10), DI computes the causal influence between variables by limiting the support of the causal variable only up to the current timestamp (equation 11). For our formulation of DI, we combine techniques from information theory and kernel-based spike train representations. Using methods shown in (*40*) for entropy estimation, we circumvent tedious probability estimations for the metric (Figure 1**B**, equations 13 - 16). Additionally, the formulation allows for DI estimation in a trial-wise manner.

Across neurons within the same cortical column, direct causal influences were identified in three steps (Figure 2**A**). First, we evaluated the DI between each pair of neurons. Next, we eliminated connections with non-significant DI by testing against the null hypothesis of non-causality (p < 0.05, *N* = 100 randomizations). To construct the surrogate data for the null hypothesis, we destroyed the causal structure between neural activity by repeatedly shuffling the spiking activity of the causal neuron. Lastly, direct influences in the neural network were teased apart from indirect influences by conditioning the DI on ‘side variables’ (equation 12). A ‘side variable’ comprises activity from a single neuron potentially involved in a cascading or proxy topological connection with the pair of neurons under study (Figure 2**A**, see Methods, spike-spike interactions). Eliminating any indirect connections is essential for neuroscientific studies, especially in the visual cortex, where neuronal networks are heavily recurrent in their connectivity. It must be noted that although DI does not determine the excitatory or inhibitory nature of the connection, it can recognize both types of influences (see Methods, Simulations, Figure S4). To aid interpretation, we use directed graphs to summarize causal connectivity patterns. Each edge and its direction in the graph is identified by aggregating the identified causal connections between the pair of nodes under study. The edge weights of the graphs are constructed to describe the strength of the connection as the averaged DI across all direct connections. For instance, an edge would be constructed between two nodes if a connection is identified between them in any one of the sessions. However, the edge width would reflect the average DI across all trials and sessions such that DI = 0 for non-significant connections.

**Figure 2:**
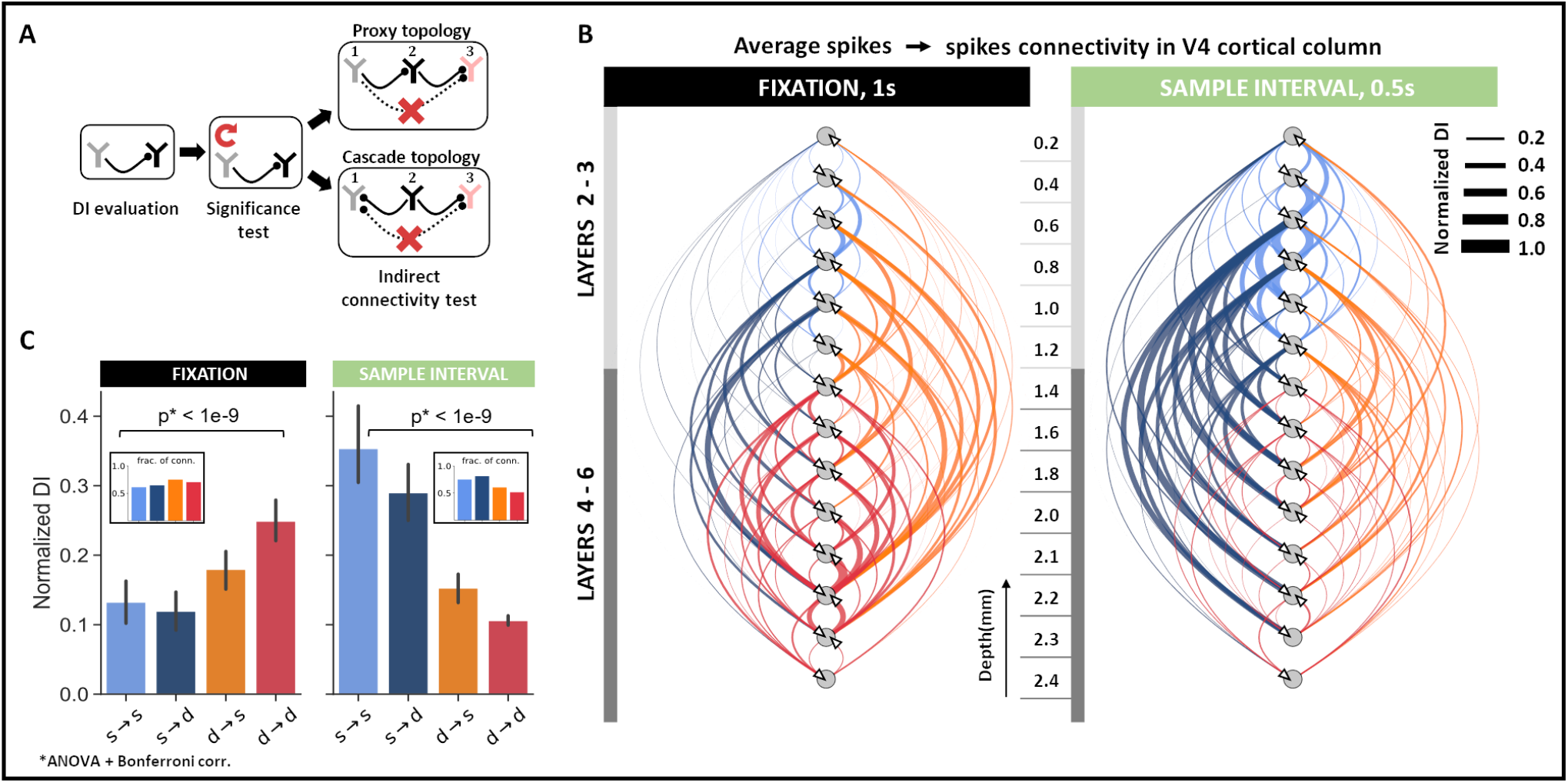
Spike–spike interactions within cortical columns of V4. **A,** Schematic to determine direct connectivity between neurons. Significant connectivities were identified against shuffled spiking in the causal neuron for *α* = 0.05. Indirect connections were removed by systematically conditioning on neurons that formed cyclic connectivities. **B**, Direct causal influences between neurons in the V4 cortical column across all sessions (N = 123 units) during fixation (1s) and early sample (0.5s) intervals. Edges are weighted by the average DI between directly connected neurons at corresponding cortical depths. **C**, Connectivity trends between superficial (s, layers 2-3) and deep (d, layers 4-6) layer neurons in the fixation and early sample intervals. Mean ± SEM across all direct connectivity.

To determine whether the neuronal interaction patterns in V4 change throughout the trial, we separately analyzed direct causal influences in the fixation and early sample interval [0-0.5]s. Across fixation and sample intervals, we observed a reversal in the within-column causal influence patterns. During fixation, the dominant pathway of causal influence included the deep → superficial and the deep → deep layer connections. The effect was two-fold. We not only observed an increased influence of deep layer neurons on the cortical column, evident via increases in DI (ANOVA, *p* < 0.001), we also noted an increased proportion of connections originating from the deep layers (Figure 2**C, B**-inset). By contrast, during the sample interval, causal neurons were concentrated mainly in the superficial layers such that there was an increase in the strength and proportion of connections originating from the superficial layer neurons (ANOVA, *p* < 0.001).

Overall, superficial layer neurons specialized more in processing information during the sample period, while deep layer neurons controlled information processing during fixation. Additionally, in line with anatomical projection patterns, these findings suggest that superficial layers support feedforward pathways channeling information from lower to higher-order cortical areas, while deep layers support feedback pathways.

### Gamma bursts are the earliest visual response in V4

Having studied interconnectivity at each scale, layer-specific patterns across the two scales became evident. During sensory stimulation, modulations in both gamma bursting and neuronal connectivity patterns were observed in the superficial layers. Further, modulations in beta bursting and neuronal interactions were confined to deep layers during fixation. To gain insights on the directionality of influence (if any) between the scales, we studied the temporal latencies of stimulus-evoked responses in both bursting activity and spiking in V4.

Using the high time resolution of the LFP point process, we noted the response latency of beta and gamma activity in each trial as the start time of the earliest burst that appeared during the 200 ms interval after stimulus presentation. The start time of each burst considers the burst length, *L* (see Methods, Burst length), and is selected as *t* = *L/*2 − *τ* seconds, where *τ* is the time-point corresponding to the center of the burst. For each of these bursts, we also identified the half-power point as the time at which the burst’s instantaneous power is closest to half the local maxima. For spiking activity, we estimated trial-wise response latencies as the time of the first spike in a 20 ms window around the neuron’s maximum response during the 200 ms interval after stimulus presentation (Figure 3**A**). Statistical analysis of the onset times across the signal modalities revealed significantly different latencies (Figure 3**C**, ANOVA, p < 1*e*^*−*3^). In most trials, the first gamma bursts began 50 ms post-stimulus presentation such that the gamma bursts were already at their half-power point 68 ms after stimulus onset, approximately 30 ms prior to the spike responses (Figure S6). In comparison, the first spiking responses appeared 93 ms post-stimulus presentation - registering an ∼ 40 ms delay between the first gamma event and spiking responses in V4. This early initiation of gamma bursts, as compared to the evoked neuronal response, could also be observed in the trial-averaged power across depths (Figure 3**B**). Lastly, beta bursts were fewer, much delayed, and were most likely to appear after spiking activity at 160 ms post-stimulus presentation. Overall, the sequence of responses followed the order: gamma, spiking, and, finally, beta activity.

**Figure 3:**
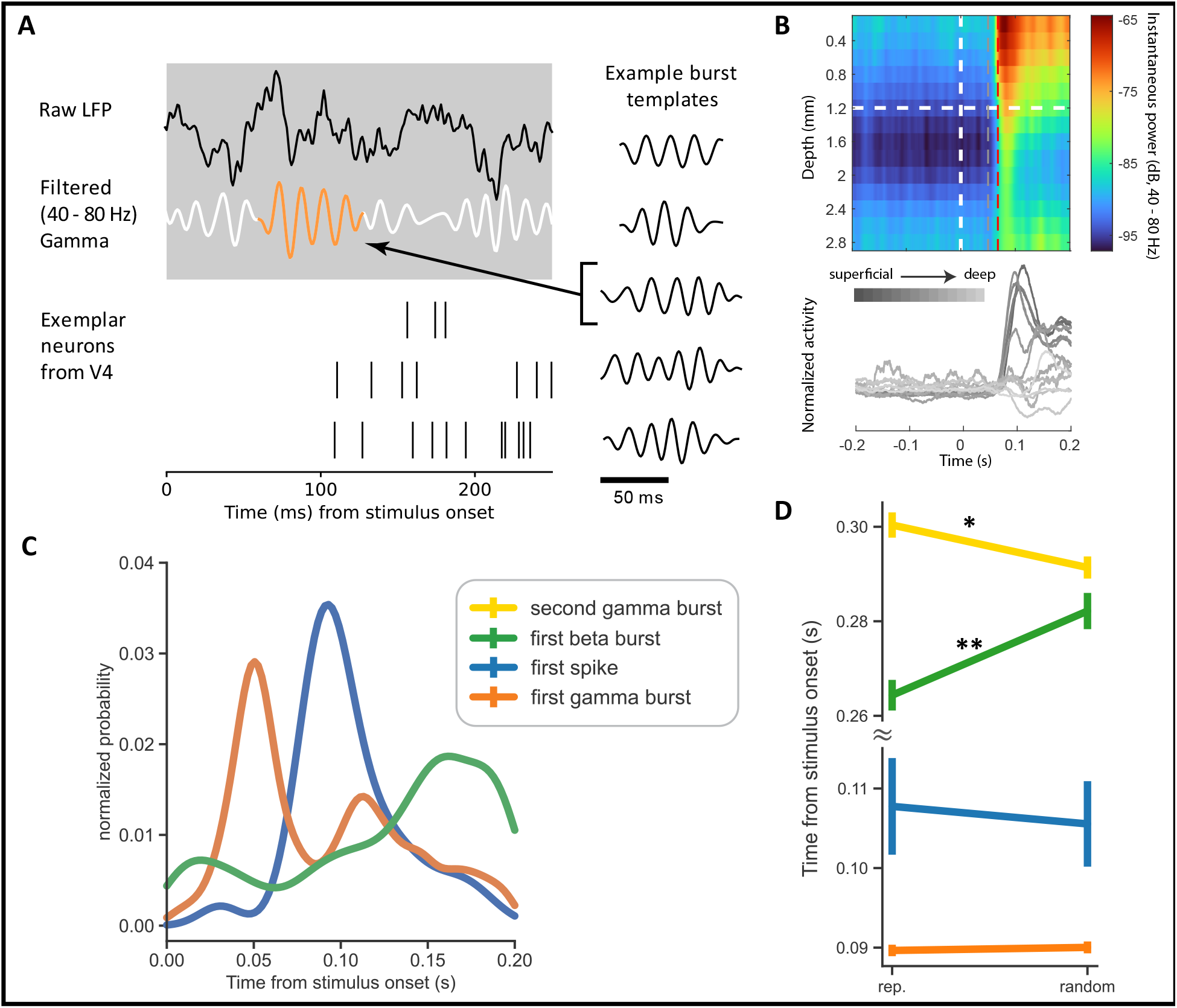
Visually evoked onset response latencies in V4 spiking and LFP activity. **A** Zoomin on neural activity after stimulus onset indicating the dynamics of raw LFPs (black), gamma oscillations (white), gamma bursting (orange) and spiking activity. Right column highlights template that maximally match the gamma burst along with other example templates. **B** Top: Trial averaged instantaneous power in the gamma frequency range of 40 - 80 Hz as a function of cortical depth. The plot also demarcates the time points associated with maximum probability of gamma burst initiation (gray dotted line) and half-power (red dotted line), respectively. Bottom: Peri-stimulus time histograms evaluated using rectangular smoothing for single neurons across the cortical depth. Exemplary plots from subject 2 and session 1.**C**, Distribution of time to first spike/burst in response to sample stimulus across all units and layers in V4. **D**, Signal response latencies separated into trials of randomized and repetitive stimuli (** *p* < 0.001). Error bars represent the 95% confidence interval.

Next, we tested if stimulus predictability impacted the onset times of bursts and spiking activity in V4 (Figure 3**D**). For this, we separately analyzed the onset latencies of spiking activity, first gamma, second gamma, and first beta event in the ‘repetitive’ and ‘randomized’ trial blocks. We identified the second gamma bursts as those that followed a gamma event in the initial 200 ms. Since the beta activity was sparse in the initial 200 ms, we relaxed the constraint on beta burst timings and considered the start times of the first beta burst over the entire sample interval. The first spiking and gamma events were strongly locked to the stimulus and were unaffected by stimulus predictability (ANOVA, gamma, p = 0.51; spiking, p = 0.3). However, the first beta burst and the second gamma burst showed significant modulations (ANOVA, p < 0.001). During repeated stimulus presentations, the first beta burst appeared ∼ 15 ms earlier than when the stimuli were randomly presented. The reverse was observed in the timings of the second gamma burst; these events appeared earlier during the randomized trial blocks by ∼ 12 ms than in the repetitive blocks.

The sequence of these stimulus-evoked responses made us wonder about the causal influence of gamma bursting on single neurons in V4. Accordingly, we set up DI to measure the layer-wise causal influence of gamma bursts on spiking activity during the early sample interval of [0 - 0.5s]. We observed a strong influence of gamma bursting on spiking that was confined mainly to the superficial layers (ANOVA, p < 0.001, Figure 4**B**). These influences were largest between superficial → superficial and superficial → deep layers (ANOVA, p < 0.001, Figure S8**A1** middle), similar to the single neuron connectivity patterns in the same interval (Figure 2**C**, right panel). Building upon the response latencies, we next analyzed the layer-wise causal influence of spiking activity on beta bursting. Spiking activity in the deep layers had a more significant causal influence on beta activity in the cortical column (ANOVA, p < 0.001, Figure S7), but the influences were generally weak. These results are in line with previous reports on oscillatory responses in the visual cortex where spike to gamma coherence was found to be most prominent in the superficial layers and spike to beta coherence in the deep layers (*2, 36, 41, 42*).

**Figure 4:**
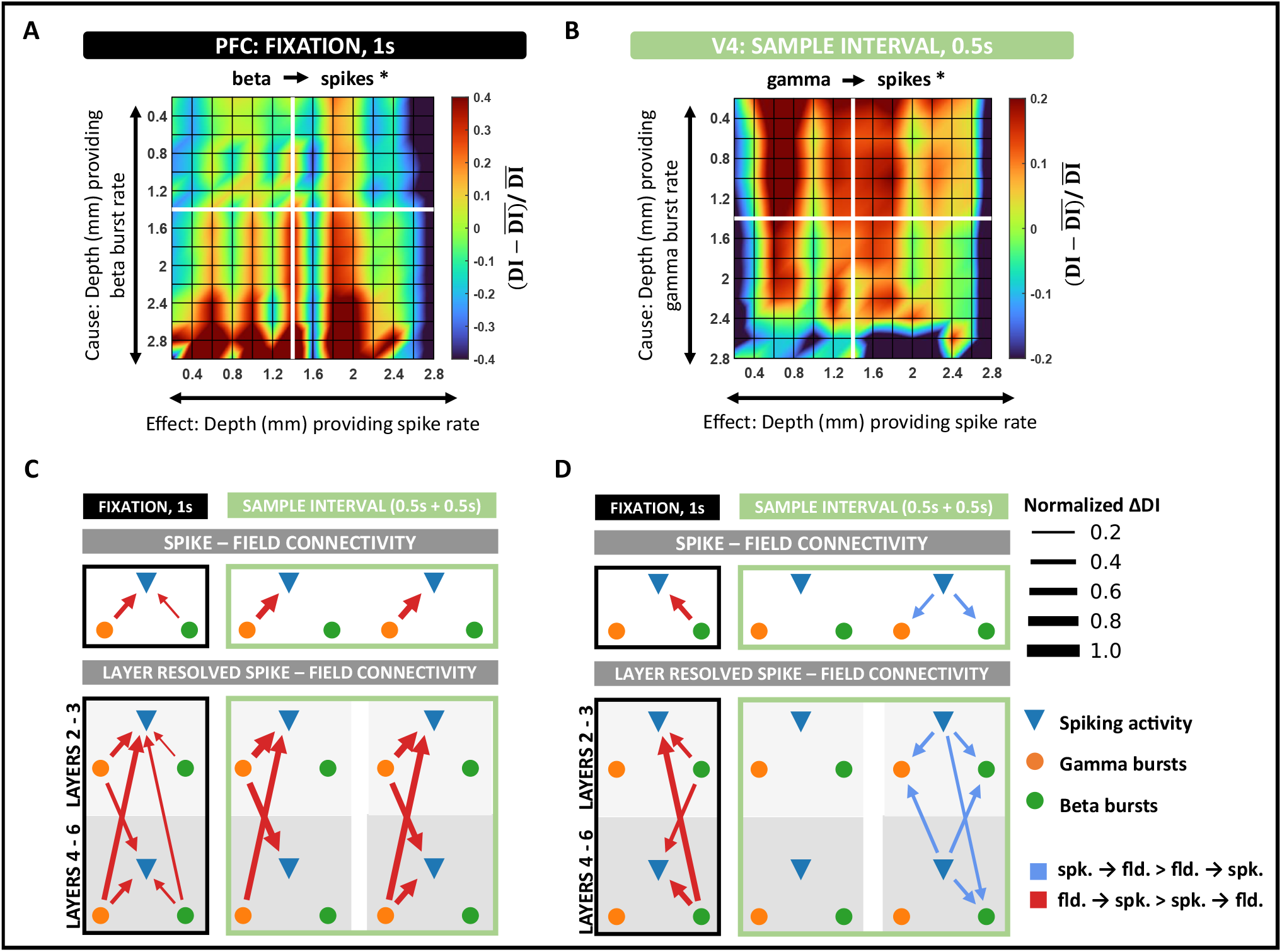
Spike–field interactions within cortical columns of V4 and PFC. **A**, Layer-specific connectivity plot in PFC from beta to spiking activity during the fixation interval (black asterisk denotes significant connection, *p* < 0.001 and Cohen d *>* 0.2). Color scale indicates the deviation of DI from the global mean DI across all layers. **B**, Layer-specific connectivity plot in V4 from gamma to spiking activity in the early sample interval (black asterisk denotes significant connection, *p* < 0.001 and Cohen d *>* 0.2). **C**, Intra – column spike – field connectivity in V4 during fixation and sample intervals. Significant connections identified as the dominant DI in opposite directions, spikes → fields and fields → spikes (*p* < 0.001 and Cohen d *>* 0.2). Edges are weighted by the absolute value of the difference in DI between opposite directions. **D**, Same as C, but in PFC.

### Spike-field interactions display consistent layer and frequency specialization but dynamic directionality across areas

Findings in the preceding section suggest a systematic causal interplay between spikes and fields in V4. To further understand these influences, we sought to study their directional asymmetries across frequencies and cortical depth. To account for the timing differences we observed in the dynamics of gamma and beta power functions (Figure S2), we separately analyzed activity in the fixation ([-1 - 0s]) and sample period ([0 - 1s]). For the same reasons, we also split the sample period into early ([0 - 0.5s]) and late ([0.5 - 1s]) intervals. Across DI estimates evaluated in individual cortical columns, the dominant causal influence between spiking activity and LFPs was identified by comparing the estimates across opposite directions (that is, *A* → *B*, if *DI*(*A* → *B*) > *DI*(*B* → *A*)). We visualized these results using directed graphs (Figure 4**C, D**), where each edge was weighted by the difference in DI between the counter-influences, ∆DI. Lastly, we color-coded the edges to represent the dominant causal variable (spikes/fields).

We found that field potentials largely drove V4 spiking activity (p < 0.001, Cohen-d *>* 0.2, Figure 4**C**), while the frequencies involved correlated with the different trial intervals. During fixation, both beta and gamma strongly influenced spiking activity, while only gamma bursts drove spiking in the sample intervals (Figure 4**C**, top row). On resolving these connections into the different cortical layers, we found interesting patterns of layer specialization in the beta and gamma connectivity patterns. Beta bursting in all layers significantly affected spiking activity in the fixation interval. However, these influences were strongest from the deep layers to the superficial layers (ANOVA, p < 0.001, Figures 4**C** column 1). The layer-specific trends of beta influence on spiking activity approximately mimicked that reported in spike-spike causal connections in the same interval (Figures 2**C** left, S8**A2** left). Across fixation and sample presentation, superficial-layer gamma activity strongly influenced spiking in the superficial layers. This influence significantly increased upon sample presentation and remained high through the sample interval (ANOVA, p < 0.001). A primary difference between interactions in the early and late sample periods was the activation of deep-layer gamma activity with a more significant influence on superficial spiking activity (ANOVA, p < 0.001).

Given the well-established role of PFC in top-down attention and working memory and its relevance to the cognitive processes engaged in the current task ((*4, 43*), 31), we examined spike-field influences in the cortical columns of PFC. While neurons in V4 were primarily driven by field potentials, causal signals in PFC switched roles between LFPs and spiking over different trial intervals. Similar to V4, beta bursts induced spiking activity in PFC during fixation. However, these causal influences were significantly higher in PFC than in V4 (ANOVA, p < 0.001). While the stimulus did not evoke any spike-field interactions in PFC during the early sample period, we observed a reversal in the directionality of influence in the late-sample period. During this interval, spiking activity induced beta and gamma bursting in the column (Figure 4**D**, top row). Intrigued by such a reversal in directionality, we wondered about the layer-wise influences of spike → field connectivity. Unsurprisingly, these influences revealed consistent frequency specialization with that of V4. Despite the reversal in directionality in the late sample period, neurons in the column drove gamma activity in the superficial layers, with the strongest influences coming from neurons in the superficial layers. While superficial layer neurons affected both gamma and beta bursting, the impact was significantly lower on beta activity (ANOVA, p < 0.001). As expected, spiking → beta influences were greatest from the deep layers (ANOVA, p < 0.001, Figure 4**D** column 3). In the fixation interval, deep beta activity strongly influenced spiking in the column (ANOVA, p < 0.001, Figure 4**A, D** column 1). During this interval, superficial beta activity also significantly modulated laminar spiking activity (ANOVA, p < 0.001); however, the influences were weaker.

### Feedforward and feedback spike-field interactions are frequency specific

To determine whether the cross-area laminar organization of spike-field communications adopts a similar specialization, we next analyzed interlaminar connectivity between V4 and PFC. Similar to the analysis in the preceding section, we identified the dominant causal influence by comparing the DI estimates across opposite directions. To aid interpretation, we constructed wheel plots to summarize the inter-area influence patterns (Figure 5). We weighted the edges by the difference in DI estimates of the opposite directions. Lastly, each edge was color-coded to correspond to the dominant directionality of causal influence, either feedforward (i.e., red, V4 → PFC) or feedback (blue, PFC → V4).

**Figure 5:**
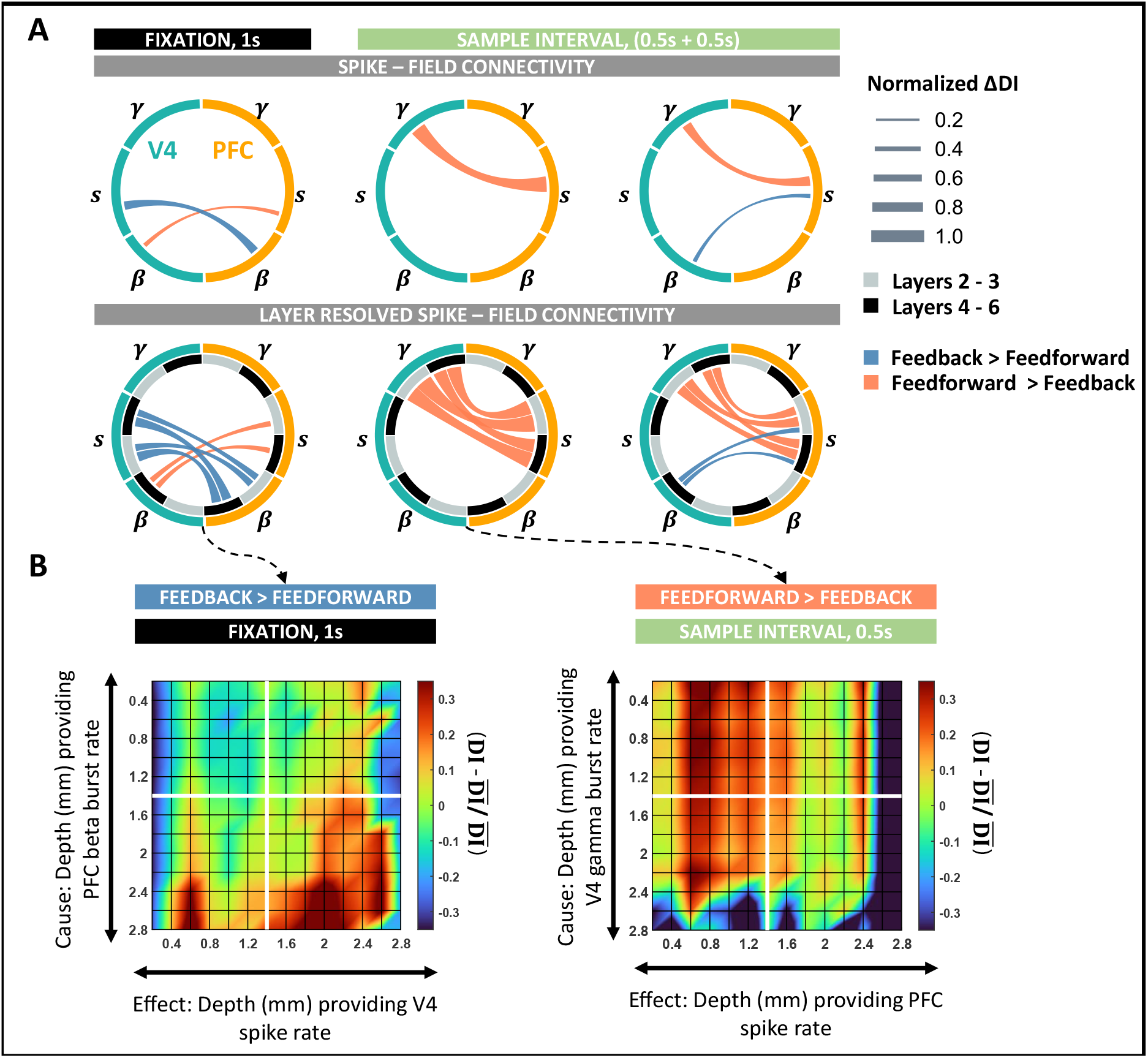
Spike–field interactions between cortical columns of V4 and PFC. **A,** (Top) Wheel plots summarizing significant feedforward and feedback spike – field connections between V4 and PFC during the fixation and sample intervals. Significant connections identified as the dominant DI in either direction of V4 → PFC and PFC → V4. (Bottom) Spike–field connectivity patterns between V4 and PFC layer resolved to deep (4-6) and superficial (2/3) interactions. **B**, Layer-specific connectivity plots between V4 and PFC for the most significant connections during fixation and sample intervals. (Left) Feedback connection from beta bursts in PFC to spiking activity in V4 during fixation. (Right) Feedforward connection from gamma bursts in V4 to spiking activity in PFC during the early sample interval. Color scale indicates the deviation of DI from the global mean DI across all layers. All significant connections were tested at *p* < 0.001 and Cohen d *>* 0.2. Edges are weighted by the absolute value of the difference in DI between opposite directions.

During fixation, overall feedback causal influences were larger than feedforward (ANOVA, p < 0.001, S9 left), where beta activity drove all influences during fixation (Figure 5**B** left, S9 left). In the feedback direction, beta activity in PFC showed causal connectivity with spiking in V4, whereas feedforward beta in V4 weakly modulated spiking in PFC (ANOVA, p < 0.001, Cohen-d *>* 0.2). Interestingly, both signals primarily originated in the deep layers of the respective areas (ANOVA, p < 0.001) and equally affected spiking in the deep and superficial layers (ANOVA; feedback, p = 0.1; feedforward, p = 0.56). During stimulus presentation, inter-laminar connections were predominantly feedforward such that gamma activity in V4 modulated spiking in PFC (Figure 5**A**, columns 2-3, ANOVA, p < 0.001, Cohen d *>* 0.2).

A layer-wise comparison of these spike-field influences revealed a strong effect of superficial gamma activity on spiking in the superficial layers of PFC (ANOVA, p < 0.001, S9**A** columns 2-3) in both the early (Figure 5**B**, right) and late sample periods. However, the strength of feedforward causal connectivity between the areas reduced over the sample interval such that stronger connections were observed in the early periods of stimulus processing. In the late sample period, feedforward gamma → spike connectivity additionally accompanied the feedback connections between spiking activity in PFC and beta bursting in V4. As per the intralaminar connections in PFC, the directionality of feedback influence was from spikes → fields such that neurons in PFC maximally modulated deep layer beta bursting in V4 (Figure 5**A**).

### Stimulus predictability modulates spike-field interaction patterns

We next investigated the spike-field asymmetries that appear in response to variations in sample predictability. Based on our results so far, we hypothesized that while repeated, predictable samples would recruit feedback pathways inducing top-down processing (PFC → V4), processing of unpredictable randomized samples would evoke signals that are fed forward up the cortical hierarchy, distilling the information in a bottom-up fashion (V4 → PFC). Further, we expected feedforward signaling to involve the gamma frequencies while feedback to involve beta frequencies. To test these hypotheses, we compared the DI estimates across the ‘repetitive’ and ‘randomized’ trial blocks in previously identified significant spike-field causal connections. We adopted a similar visualization style to the previous sections (Figure 6). We weighted the edges by ∆DI, corresponding to the difference between the DI estimates in ‘randomized’ and ‘repetitive’ trial blocks. While we maintained the directionality of influence from the previous sections, each edge is color-coded to represent the stimulus type (either ‘randomized’ or ‘repetitive’) of higher DI for that specific connection.

**Figure 6:**
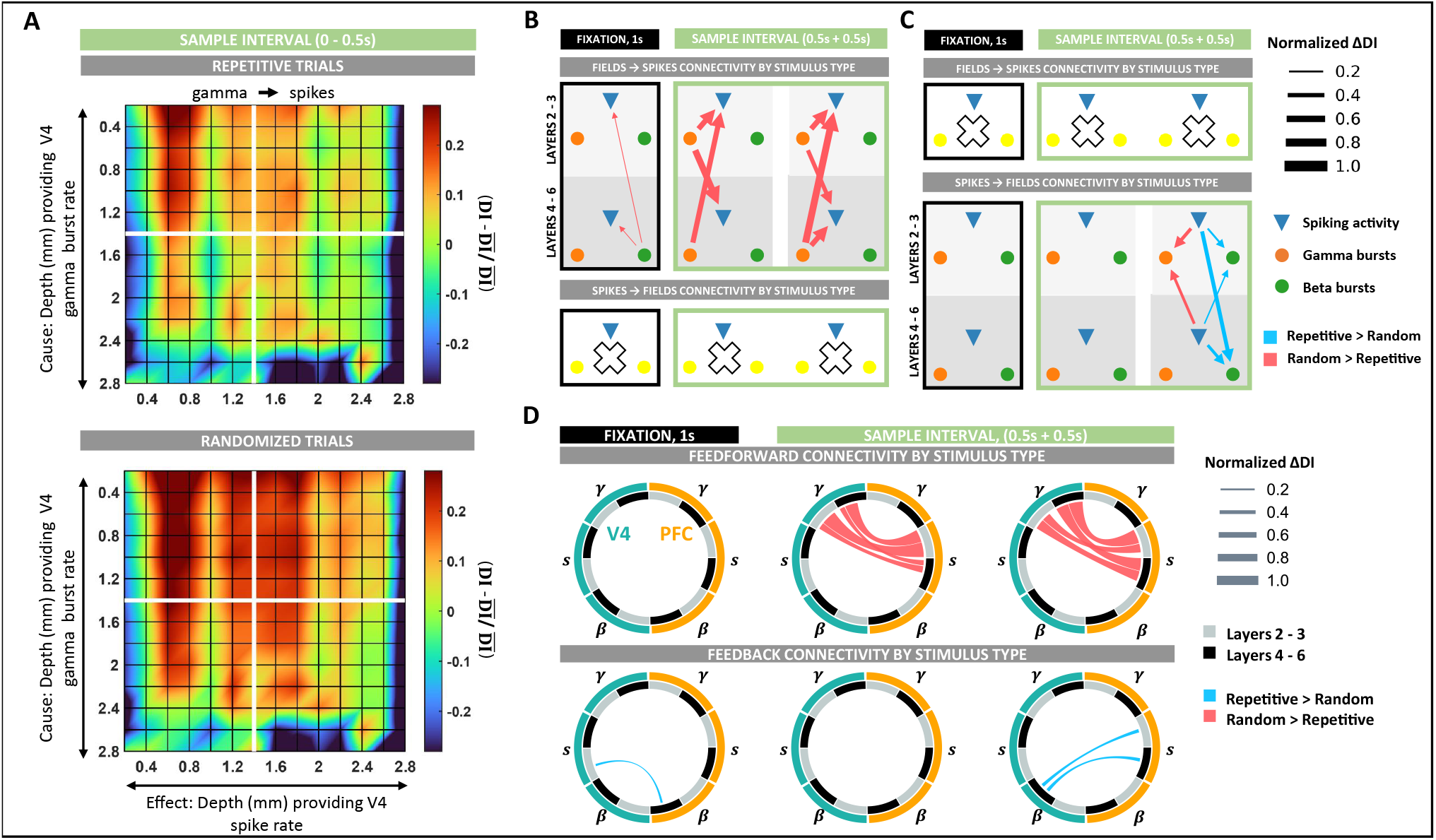
Spike–field interaction patterns modulations by stimulus type. **A,** Intra-column spike–field connectivity as modulated by stimulus predictability (randomized vs. repetitive) during fixation and sample intervals in V4 (top) and PFC (bottom). **B**, Spike–field connectivity between columns of V4 as modulated by stimulus predictability during fixation and sample intervals. All significant changes in DI between the trial groups of randomized and repetitive stimuli were tested at *p* < 0.01. Large white crosses denote an absence of significant modulations. Edges are weighted by the absolute value of the difference in DI between the two trial groups. **C**, Same as B, but in PFC. **D**, (Top) Wheel plots summarizing feedforward spike – field connections between V4 and PFC significantly modulated by sample predictability during the fixation and sample intervals. (Bottom) Same as top panel, but summarizing feedback connectivity significantly modulated by sample predictability.

First, we analyzed the variations in intralaminar spike-field connectivity in V4 and PFC. In V4, spike-field connectivity was stronger during ‘randomized’ stimulus presentations (Figure 6**B**, S10**A**). During this trial block, intralaminar connectivity between gamma → spike increased in the early sample interval such that the most significant increments were between superficial layer gamma to deep layer neurons (ANOVA, p < 0.001, 6**A**, 6**B** middle, S10**A** middle). Next, in the late sample period, we observed the most significant enhancement in connectivity from deep gamma → superficial spiking activity (ANOVA, p < 0.001, 6**B** right, S10**A** right). Lastly, during fixation, the effect of deep layer beta activity on spiking was weakly higher to ‘randomized’ stimuli (6**B** left, S10**A** left).

In PFC, modulations in intra-laminar connectivity to sample predictability were only observed in the late sample period and only in the connections from spike→field (6**C** right, S10**B**). These influences had a dual response to stimulus type. While spike → beta influences were enhanced during repeated stimulus presentations, spike → gamma influences were stronger during the ‘randomized’ trial block. When resolved layer-wise, neuronal connectivity was higher to superficial-layer gamma during randomized stimuli (ANOVA, p < 0.001) and to deep-layer beta during repeated stimulus presentations (ANOVA, p < 0.001).

As expected, variation in intralaminar connectivity to sample predictability implicated a role of gamma bursts in the ‘randomized’ trial blocks and beta bursts in the ‘repetitive’ trial blocks. To determine if these variations also associate with feedforward and feedback processing, we applied our analyses to study the connectivity across areas in the two trial blocks. Indeed, interarea connections between V4 and PFC showed a clear separation in connectivity between topdown and bottom-up processing (Figure 6**D**). Whereas feedforward connections reported significantly larger causal influences while processing unpredictable ‘randomized’ stimuli, all feedback connectivity was enhanced when the stimulus was predictable (‘repetitive’ trial blocks). In the sample period, feedforward connections between gamma in V4 to spiking in PFC was most significantly enhanced. While in the early sample period, we observed the most significant increases along the pathways from superficial layers in V4 to superficial layers in PFC (ANOVA, p < 0.001, Figures 6**D** top-middle, S11**A** middle), in the late sample period, this increase was strongest along the pathways originating in the deep layers of V4 to the superficial layers in PFC (ANOVA, p < 0.001, Figures 6**D** top-right, S11**A** right). Lastly, feedforward connections during fixation did not display any modulation to stimulus predictability.

Along the feedback pathway, spike-field connections strengthened when the sample was predictable. During fixation, repeated stimulus presentations enhanced connectivity between beta bursting in PFC and spiking activity in V4. Specifically, these connections showed increased connectivity from deep layers in PFC to superficial layers in V4 (ANOVA, p < 0.05, Figures 6**D** bottom-left, S11**B** left). While we registered no feedback connections during the early sample period (Figures 6**D** bottom-middle, S11**B** middle), the influence of spiking activity in PFC on beta bursting in V4 increased during the late sample period (Figures 6**D** bottomright, S11**B** right). Like connectivity modulations during fixation, these connections showed the largest modulations along the deep to superficial pathways (ANOVA, p < 0.05).

## Discussion

In this work, we studied the causal dynamics of spike-field interactions by relating spiking activity with oscillatory bursts in the LFPs. Using a novel unifying point process representation for the multiscale brain activity, we systematically analyzed spike-field connectivity within and across the visual and prefrontal areas. We independently analyzed the spike-field causal flow as the brain switches between two modes of information processing: 1) the bottom-up mode during which novel sensory inputs had to be processed on every trial, and 2) the top-down mode when the same stimulus was presented on every trial alleviating the need to process it. Spiking activity and LFPs are analyzed in the same Hilbert space using the same methodologies and time resolutions. We also account for the distinct dynamics of neural activity in different cortical layers, task rules, and stimulus types. We find that spike-field communication is more complex than previously assumed (*44–46*) in that it is a dynamic task-modulated relation where field fluctuations do not always reflect the temporal pattern of spiking activity near the electrodes. Spike-field connectivity also adheres to a consistent laminar-specific structure over the cortex, similar to hierarchical anatomical connections. Specifically, our results reveal two separate communication pathways during top-down and bottom-up information processing: 1) interactions between gamma and spiking that traverse the pathways originating in superficial layers and travel towards superficial layers during behavior that induces bottom-up processing, 2) interactions between beta and spiking activity that originate in the deep layers and travel towards the deep layers during behavior that induces top-down processing. While our results are consistent with previous findings on frequency-specificity (*2, 36, 43*), we also demonstrate the dynamic change in interaction patterns with high time resolution. Below we summarize our observations.

### LFPs and spike generation

Rooted in the biophysical explanation of LFPs, it is commonly accepted that the oscillatory structures observed ubiquitously in the signal reflect synchronized inputs to a given area (*47, 48*). However, it is unclear if LFPs are a mere reflection of synchronization due to an underlying rate modulation or if these oscillations also provide a framework that allows for precisely coordinated spiking as predicted by an active assembly (*49–51*). Our results provide support for the latter hypothesis. Firstly, the stimulus-related spiking response of single neurons in V4 occurs about mid-way through the first gamma burst in the same area (Figure 3). This observation is in line with neural synchronization models that suggest that optimal communication between two groups of neurons can be achieved when synchronized activity of presynaptic neurons, arriving as bursts at the post-synaptic cell, allow for maximal input density at the target neuron; thereby, eliciting a response (*52, 53*). In V4, this mechanism appears to be implemented by gamma bursts during the early sample period. We speculate that the registered gamma bursts in V4 are not local to the cortical column but may be outputs from presynaptic pyramidal neurons from an upstream area (like V1) or surrounding columns in the visual cortex (*54–56*). This result also aligns with response latencies reported in (*56*), where engagement of V1 neurons begins around the same time we observed gamma bursts starting to form in V4. The direct effect of gamma bursting on spiking in V4 is later captured as gamma → spike influence in our causality analysis Figure 4**C**. Secondly, layer organization of spike-spike connectivity in V4 mimicked the organization of field → spike interactions in the column corresponding to either frequency range (beta during fixation, gamma during sample presentation). These associations support the active engagement of LFPs in coordinating spiking activity. Lastly, 75% of all the significant spike-field causal connectivities identified in the study were directed from fields → spikes, suggesting that oscillatory bursts in the LFPs carry information that significantly influences future spike generation.

### Frequency specificity of spike-field interactions

Properties of oscillations in the beta and gamma frequency ranges have implicated their role in gating and control. While ubiquitously observed over the cortex, they are anti-correlated (*57, 58*), Figure 1**E**. In the visual cortex, gamma power is high and beta low during sensory stimulation, while the reverse is true when a stimulus is filtered, ignored, or absent (*36, 59, 60*). In PFC, the same frequencies have been implicated in the encoding, maintenance, and read-out of working memory signals (*37, 58, 61*). These findings have led to the general idea that top-down signals are fed back through beta, and gamma helps maintain the spiking activity carrying sensory inputs, thereby aiding feedforward communication.

In our studies, these specializations broadly translate to spike-field causal connectivity. Generally, connectivity between spiking and beta activity was observed during fixation and late sample periods, i.e., during the absence and filtering of sensory information. Contrary to beta, connectivity between gamma and spiking was strongest when the subjects were visually stimulated. Local spike-field interactions complemented global interactions showing a similar task-modulated pattern. However, in the late sample period, we found that beta and gamma pathways were simultaneously active over local and long-range connections. During this period, gamma bursting in V4 maintained a feedforward influence on spiking activity in PFC, which simultaneously controlled gamma and beta bursting in the PFC column. In the same interval, feedback influences were noted from spiking in PFC that stretched over beta bursting in V4. Such dual processing in the beta and gamma pathways implicate a complex system that enables simultaneous local information integration and feedback pathway preparations between higher- and lower-order areas (*62*). Notably, this dual processing by PFC followed after a period of quiescence. Although V4 gamma maintained a feedforward influence on spiking in PFC throughout the sample period, information processing in PFC was only noted in the latter half. These timing delays are within the previously observed ranges for cortico-cortical interactions (*63*) and represent an activation sequence of critical components of the attentional-control brain network. Our findings suggest a signaling system where distinct high and low-frequency signals mediate feedforward and feedback processing, respectively. Herein the top-down effect of beta bursting helps regulate the processing of bottom-up inputs served by gamma bursts and spiking activity.

### Hierarchical organization of spike-field connectivity

Current dogma holds that the cortex is hierarchically organized, where cortical areas are classified by their laminar origin and termination patterns (*64–66*). In this view, feedforward connections originate mainly from superficial pyramidal cells and target the middle layers. Feedback connections originate primarily from deep pyramidal cells and target layers other than the middle layers. While the current description is a simplified depiction of the complex brain networks, recent studies have revealed physiological distinctions in layer-wise dynamics. For instance, neuronal synchronization is asymmetrically distributed across cortical layers, wherein superficial layers show higher synchronization and spike-field coherence in the gamma frequencies, while the deep layers prefer low frequencies (*2, 36, 41, 67*). Granger-causality studies further support this functional asymmetry, discernable even over long-range, inter-areal connections (*3, 68*). Given ample evidence supporting the role of gamma (resp. beta) in feedforward (resp. feedback) processing, it is likely that superficial and deep layers constitute functionally distinct processing streams.

We noted the first evidence of the functional separation between layers in the direct causal connectivities of single neurons in the V4 cortical column. While during fixation, we observed more recurrent and lateral connections within the deep layers, the network connections completely reorganized on visual stimulation, predominantly recruiting superficial layer neurons for stimulus processing (Figure 2). This result strongly supports that superficial layers are placed hierarchically lower than deep layers and support feedforward processing. Next, the dynamic interplay between the low and high-frequency spike-field causal influences revealed a further distinction between layers. These dynamics were complementary not just in their timings but also in their layer-specific influences. While superficial-layer gamma most strongly influenced spiking in the same layers during feedforward processing, deep-layer beta selectively affected spiking in the deep layers while controlling feedback processing. This layer-specificity of neural rhythms was held even when the directionality of influence reversed in the PFC in the late sample period. Alongside asymmetries in the within-column spike-field causalities, the specializations were consistent over long-range connections between V4 and PFC (Figures 4, 5). Lastly, in the context of hierarchy, it is worth noting that the directionality of influence from spikes → fields was only observed in the PFC in the late sample period. While further systematic studies on PFC connectivity with other intermediate areas are required to comprehend this observation fully, these results are promising because of the role of PFC as a source for topdown signals that biases selection in early visual areas in favor of the attended features (*69*).

### Relation with predictive coding models

Predictive coding models suggest a gating mechanism, wherein experience generates predictions that attenuate the feeding forward of predicted stimuli while only projecting the residual errors of prediction forward. These unpredicted errors act as updations that correct the internal prediction generation model. While theories on model implementation are varied (*65,70–72*), our results are most consistent with a neurophysiological explanation based on distinct neuronal rhythms (*65, 73*). These models suggest that superficial gamma activity is responsible for the feedforward projection of prediction errors, and predictions are fed back to the deep layers via a beta channel. In this framework, predictions prepare the beta channels that actively inhibit the input-processing gamma pathways in the sensory cortex, eventually reducing the feedforward outputs. Therefore, prediction errors result from a lack of inhibition of the feedforward pathways by low-frequency top-down signals rather than from specialized circuits finalizing comparisons between the prediction and inputs.

The unique setup of the current task incorporates modulation of the stimulus predictability in alternating blocks. Since all stimuli were presented at full contrast, and both block types required the subjects to attend to all stimuli, prediction errors are less likely to be modulated by variations in sensory evidence or attentional modulation. The general idea is that unpredictable randomized stimulus presentations would increase prediction errors corresponding to more mismatched inputs and internal predictions. In contrast, repeated stimuli would require a memory recall and induce fewer prediction errors. In such a case, block-wise analyses of spike-field connectivity revealed frequency-specific associations with stimulus predictability, garnering further evidence for the role of neural oscillations in directing the information flow of prediction signals (Figure 6). Generally, randomized stimulus presentations induced stronger feedforward connectivity between spikes and gamma, while repetitive presentations strengthened the feedback associations between beta and spiking. Across-layer influences were also affected by modulations in stimulus predictability across local and global networks. However, no specific generalization between layers could be drawn.

### Deviations from common generalizations

Not all connectivity between LFP frequencies and spiking strictly adhered to all the above generalizations. Although relatively weak, gamma → spiking influences were significant in the V4 columns even in the absence of any visual stimulation (during fixation, Figure 4**C**). Interestingly, this influence of gamma on spiking activity was greater from the deep layers. A similar increase in deep gamma → superficial spiking was also observed in the late sample period in V4. Second, contrary to our expectation, we observed feedforward beta → spiking influences between V4 and PFC during fixation (Figure 5). These V4 beta bursts also reported increased influence on local spiking activity during the randomized trial blocks (Figure 6). These deviations suggest that LFP frequencies assume dynamically changing roles involved in network activity that are more complex than simple generalizations.

### Improvements over other causal inference methods

In computational neuroscience, the assessment of multiscale causality presents a challenging computational problem because of the widely disparate statistics of spiking and field potentials. Recently, Granger causality measures have been adapted to study spike-field causality (*44, 74–77*). While these models were some of the first to achieve causal inference on hybrid neural signals, they are constrained by important limitations. Firstly, all the proposed models assume that the true data-generating process and the corresponding causal effects of the variables on each other are linear, whereas neural activity is a non-linear process. In such a case, the interpretation of the connectivity estimates can be misleading (S4 **F**). Our Hilbert space approach for DI estimation extends the linear model mathematics to non-linear modeling in the input space, where the reproducing kernel Hilbert space can handle different types of nonlinearities. Second, the timescales of changes in singleunit activity and LFPs vary vastly. Naturally, the fastest time scale should set the limits on the diagnostic model, i.e., inferences must operate on signals with millisecond resolution. Studies that derive support from Fourier analysis methods inevitably fail to capture such high-resolution representations (*78*) due to the trade-offs between time and frequency resolutions (*44, 74, 75*). Our causal estimation framework operates on a high time-resolution construct of LFPs based on bursting activity in specific frequency bands, thereby maintaining time resolutions across the signal modalities.

While the model recognizes critical causal events, the inference is limited in its time resolution, wherein the entropy estimator requires enough samples (300 - 500, (*40*)) to avoid improper estimations. Third, studies that adopt a modeling approach for causal analyses rely on ‘plug-in’ estimators where the data is first fit to a model distribution and then plugged into the causality estimation framework (*76, 77*). Although plug-in estimators are intuitive, estimating the distribution is a complex problem, both parametrically and non-parametrically. While parametric estimators struggle to balance tractability and oversimplification, non-parametric techniques are computationally demanding and susceptible to overfitting. In either case, making an i.i.d assumption on the data is inappropriate for studying causal analysis. Our causal estimation framework does not assume prior information on signal distribution or independence in either modality, making it a data-driven approach. Lastly, most studies on spike-field analysis do not address the non-causal effects of spike-related transients in the LFPs (*20*), thereby leading to spurious causal estimations. In our studies, we incorporate a rigorous characterization of bursting activity in terms of burst structure, signal-to-noise ratio, transiency, and frequency specialization, alleviating the impact of spike-related transients in LFPs. Overall, our approach identifies functional relations between spikes and fields in a neurophysiologically sound and computationally appropriate manner.

Here, we also emphasize that in the context of brain connectivity patterns, it is generally impossible to distinguish absolute direct influences from indirect influences if not all nodes of the functionally connected network are sampled. While the problem is not theoretical, it has important implications for interpreting neuronal interactions (*79, 80*). Under the circumstances, care must be taken when making causal inferences. Nevertheless, with increased coverage of the brain areas, methods such as ours and those by others will be crucial in inferring the direct and indirect pathways of brain signaling. Lastly, while the current study only focuses on spike-field interactions in the cortex, it only partially represents all connectivities plausible in the neural substrate. The current framework can seamlessly translate causal relations between single neurons, LFPs, and cross-frequency relations in the LFPs. Future studies will consider these analyses while incorporating signals from intermediate areas such as frontal eye fields and posterior parietal areas. These examinations will merit insights into the area-specific properties of spike-field relations and the activation sequence of an attention-driven brain network.

## Methods

### Experimental Setup

The data analyzed in this study comprised of multi-laminar recordings from the visual area 4 (V4) and prefrontal cortex (PFC) of macaque monkeys (*Macaca mulatta*) trained to perform a delayed match to sample (DMS) task (*2*). We analyzed ten-session recordings from two monkeys that performed five sessions each. Subjects were implanted with linear array U and V probes (from Plexon) using a custom-machined carbon PEEK chamber system with recording wells placed over the visual/temporal and frontal cortex. The center of each chamber was overlaid with the primary recording area of interest and optimized for an appropriate angle for perpendicular recordings relative to the cortical folding. The number of laminar probes varied between 1 and 3 for each session and brain area. Each probe comprised 16 electrodes with an intersite spacing of 100 or 200*μ*m, culminating in a total linear sampling of 3.0 to 3.1 mm on each probe. Channels from the top of the cortex to a depth of 1.2 mm were classified as superficial layer channels, and deep layer channels were those below 1.2 mm. All surgical and animal care procedures were approved by the Massachusetts Institute of Technology (MIT)’s Committee on Animal Care and were conducted per the guidelines of the National Institute of Health and MIT’s Department of Comparative Medicine.

The experiments were conducted inside a sound-proof behavioral testing booth comprising a primate chair placed 50cm away from an LCD monitor (ASUS, Taiwan). Subjects remained seated throughout the session and were trained to perform the DMS task using positive reinforcement. Monkeys that fixated on a point in the center of the LCD screen for a complete duration of 1s were presented with one of three cue objects. The object remained on the screen for 1s and disappeared during the following delay period. In experiments with monkey-2, the delay period lasted for a uniformly picked time interval between 0.5 − 1.2s, whereas in experiments with monkey-1, the delay period was a fixed time interval of 1s. At the end of the delay period, a search array consisting of the cued item and one or two confounding objects appeared on the LCD screen, each occupying a different visual quadrant. The distractor and its position were randomly chosen. A successful trial entailed the subject performing a saccade towards the cued object, for which they received a few drops of diluted juice for positive reinforcement. The trial was terminated if the subject broke fixation at any time. Finally, the cued object’s predictability was manipulated with repetitive or randomized cueing. During randomized cueing, the cue was randomly sampled from 3 objects and presented in each trial. Contrarily, repetitive cueing consisted of the same cue presented in each trial. Each block lasted for 50 trials, where the initial cueing type was randomly chosen. The trial schematic is presented in Fig.1**C**. For specific details on behavioral performance, please see (*2*)).

Throughout the session, neural activity was recorded with Blackrock headstages (Blackrock Cereplex). The signals were sampled at 30 kHz and bandpass filtered between 0.3 Hz and 7.5 kHz using a first-order Butterworth filter. We used a low-pass Butterworth filter with a cutoff frequency of 250 Hz and sampled at 1 kHz to extract the local field potentials. A Plexon offline sorter was used to perform spike sorting manually. For analyses, both field potentials and spiking activity were downsampled to 500Hz.

### Generative Model for Oscillatory Events

We used a generative model to detect oscillatory bursts from LFPs. Here, we briefly describe the method; for further details of the model, refer to (*37*). Oscillatory bursts are brought about by external inputs that synchronize neural activity in the ‘spontaneous’ state of the neural dynamics. Spontaneous state activity follows a 1/f power law with amplitudes that approximate a Gaussian distribution. Oscillatory bursts, on the other hand, deviate from this ‘Normality’ and appear as a high-amplitude rhythmic activity that wax and wanes (*81*). The generative model for oscillatory events builds on this two-state hypothesis. The model simplifies LFP modeling by operating on established cortical rhythms of interest in parallel (*82*). Specifically, the model defines single-channel, bandpass filtered traces of neural recordings, 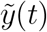, as a linear combination of spontaneous activity, *n*_0_(*t*) and oscillatory bursts, *ŷ* (*t*), as shown in equation (1). Next, each oscillatory burst is modeled as the impulse response of a linear filter as shown in equation (2) (*83*). Each filter represents a ‘typical’ temporal response of population-level synchronization in the frequency range under study. The set of filters define a dictionary of templates, 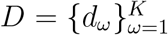. Specifically, each oscillatory burst is represented as a shifted, scaled version of a filter, *d*_*ω*_, such that 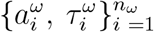 are the amplitude scalings and the timestamps of the oscillatory events represented by the learned filter (equation (2)).

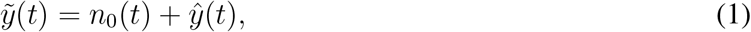

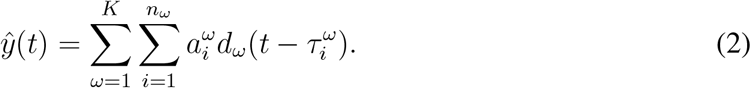

The features of each burst, namely, its time of occurrence (*τ*_*i*_), maximum amplitude (*α*_*i*_), specific frequency (*ω*), and duration (*δ*_*i*_) then constitute a Marked Point Process (MPP); wherein lies the model’s advantage to characterize the properties of the oscillatory events in a high – resolution representation of channel activity.

The generative model realizes this (marked) point process representation in two phases: 1) a learning phase for estimating the dictionary, *D*, from the training data; 2) a testing phase for constructing the point process from the test signals, given the dictionary.

As a first step in the training phase, we delineate the search space for oscillatory bursts using a step called ‘denoising”. ‘Denoising’ involves discriminating between the spontaneous activity, *n*_0_, and the oscillatory bursts, *ŷ*, using a correntropy-based clustering technique (*84*). The method locally differentiates between the two using their complete probability density information. In the final denoising step, a threshold, *κ*, is evaluated as the minimum norm among the putative oscillatory events. After denoising, learning is implemented by alternating optimizations between sparse coding and (correntropy-based) dictionary learning, paralleling a K–means clustering algorithm to learn the oscillatory filters in a data-driven fashion. The sparse-coding stage implements the learned threshold to identify bursts from spontaneous activity. The training of the K-means clustering model is terminated as a specific number of iterations is reached or when an upper bound on the Frobenius difference norm between successively determined dictionaries is met.

Finally, constructing the MPP from the test signals is done in a straightforward and computationally efficient way. Each filter in the learned dictionary is convolved in parallel with each channel from the test data. Following this, *κ* is adapted to the test data, and thresholding isolates the oscillatory burst events from spontaneous activity in the test data. In this way, the generative model can characterize the properties of each oscillatory burst and enable an easily interpretable point process representation of multi-channel neural data. (Figure 1**A**). Overall, the model derives support from two user-set hyperparameters: 1) the maximum duration of an oscillatory burst event, *M*, and 2) the number of filters, *K*, in the dictionary. We link the MATLAB implementation of the model here: https://github.com/shailajaAkella/MPP-EEG_Ver2.

#### Burst length

The duration of each oscillatory burst is determined to eliminate the non-oscillatory parts of the identified burst. First, a smoothed envelope of the detected event is estimated using Hilbert Transform and a moving average filter. Following this, two points of minima on the envelope that are separated by at least *M/*2 samples are identified. Finally, the signal length between the two minima determines the duration of each burst. These steps are summarized in (3 - 6).

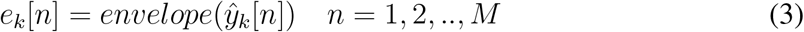

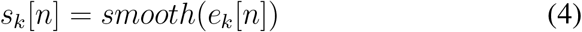

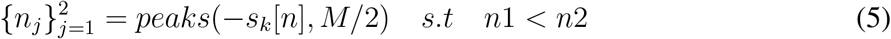

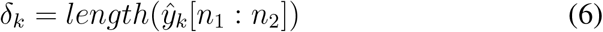

#### Burst power functions

The burst power (*85*) is an extension of the classical intensity function of point processes, wherein the intensity function is weighted by the transient power of the event at its time of occurrence. Given the timings of each oscillatory event, *τ*_*k*_ : *k* = 1, 2, …*N*, the construction of the burst power function is summarized in equations (7–9). First, transient oscillatory power, 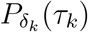, is estimated as the norm-square of the signal amplitude values in the duration of each oscillatory burst (equation 7). Next, the event trains are weighted by their local power, 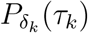, to form the marked event density, 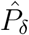 (equation 8). Finally, an estimate of the local burst power is obtained via Gaussian kernel smoothing (equation 9).

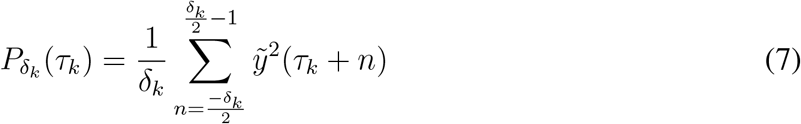

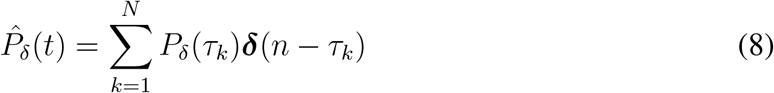

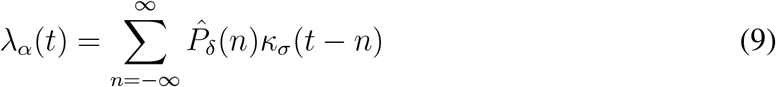

#### Parameter selection

The generative model is applied to learn the typical templates of burst patterns from (bandpass) filtered traces of LFP recordings in the broad beta (8 - 30 Hz) and gamma (40 - 80 Hz) frequency ranges. Learning on the model was performed per session, area, and frequency range, culminating in 4 dictionaries per session (two areas and two frequency ranges). We perform zero-phase FIR filtering to obtain these signals, while model training is performed on a single channel (channel at 200 mm depth) which we then exclude from the testing phase. The designed bandpass FIR filters have the following properties: 1) for the beta band, the center frequency is set at 20 Hz, and the filter order is 25, 2) for the gamma band, the center frequency is chosen to be 60 Hz which culminates in a filter of order 11. We deliberately chose low filter orders to avoid ringing in the filter response. The maximum duration of each event, *M*, in the beta and gamma frequency ranges, are correspondingly set equal to 500 and 100 ms, while the number of basis vectors (dictionary atoms), *K*, is upper bound to 30 for both. We employ Silverman’s rule of thumb for bandwidth size to set all kernel bandwidths for the denoising phase (*86*).

Finally, the burst power and rate functions are estimated from a point process representation using Gaussian kernels, and the selection of the bandwidth size must be neurophysiologically informed. For this, we chose a bandwidth size that approximately covers the duration of the burst event, translating to 50 and 30 ms for beta and gamma bursts, respectively.

#### Validation

The generative model has been previously empirically validated; please refer to (*37*) for details about the validation. This section tests the model against simulated LFP data and presents a comparative analysis with the most common burst detection methodology, hard-thresholding (*58*). Simulated LFPs were generated by superimposing power law noise (*α* = 2, (*87*), Figure S3**A**) with oscillatory bursts in the 50 − 70 Hz frequency range. Each trial was about 2s long, and 100 trials were simulated, with 0-5 bursts in each trial (following a Uniform distribution). The length of each burst varied between [100 − 120] ms, and consecutive bursts were introduced at least 100 ms apart. Lastly, we also varied the signal-to-noise (SNR) of each burst between 0.4 - 0.5, such that the total SNR in each trial came up to ∼ 0.5.

Next, we set up the generative model for burst detection. We filtered the LFP signals using a broadband gamma filter between 40 − 80 Hz for model training. We set the maximum length of each burst, *M* = 120 ms, and varied the number of filters, *K*, between [10 - 100] to test model performance. For model selection and performance evaluation, we performed a five-fold cross-validation analysis of the data. Model performance was evaluated using two metrics: 1) the rate of bursts correctly estimated or true-positive rate (TPR), 2) the proportion of time points cataloged as bursts when they are not, or false-positive rates (FPR). Lastly, we compared our model’s performance against the time-frequency methods used in (*58*) for burst detection. Specifically, in (*58*), the authors identified oscillatory bursts as periods during which the spectral power surpassed the threshold, established as two standard deviations (SD) higher than the mean value for the given frequency range of that specific trial. The duration of each burst was further constrained to be at least three cycles. Similar to these methods, we used a multi-taper approach to obtain single-trial spectrograms for the LFPs (*88*). We further varied the threshold from 0.5 × SD to 3 × SD over the mean power to detect bursts while maintaining a three-cycle limit on the burst length.

The performance of the generative model was relatively stable across different dictionary sizes (Figure S3**C-D**, *SD*_*T PR*_ = 0.6%, *SD*_*F PR*_ = 0.04%). Nevertheless, the model reported its best performance at K = 50, with a TPR of 93.7 ± 2.5% and an FPR of 3.8 ± 0.4% (Figure S3**E - F**). In comparison, at a threshold of 2× SD (as implemented in (*58*)), the performance of the TF method was much poor: TPR = 24.7% and FPR = 1.1%. The method’s best TPR was observed at a threshold of 0.5× SD at 85.4%. However, a lower threshold also increased the number of false positives, and FPR rose to 12.8%. Overall, the generative model demonstrated superior performance. These results highlight the advantage of a data-driven approach for burst detection where hard-thresholding mechanisms cannot capture the variability in neuronal oscillations. These methods impose assumptions of stationarity and ergodicity on an inherently stochastic neural signal, thereby confounding the boundaries between transient bursts and background activity. Hard thresholding is generally sensitive to signal SNR, and the disadvantage of poor time and frequency resolutions makes it difficult to extract this metric and estimate precisely in time when it exceeds the baseline values. In such a case, the generative model provides an unsupervised, adaptive framework that does compromise between the temporal and spectral resolutions of neural data.

### Directed Information Estimation

In information theory, the mutual information (MI) between two processes, *X*, and *Y*, communicating over a channel with feedback, can be decomposed into two components: one component captures a causal, feedforward or *directed information* (DI) flow, and another that captures a non-causal or a feedback information flow (equation 10) (*89*). Therefore, inferences based on MI can be detrimental in brain networks wherein the neural substrate implements distributed recurrent communications.

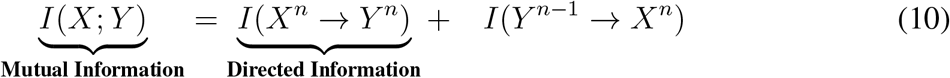

A formal definition of directed information between two stochastic processes, *X* and *Y*, is shown in equation (11), where *X*^*n*^ denotes a segment of the successive samples of the process, *X*, i.e., *X*^*n*^ = [*X*_1_, *X*_2_, …, *X*_*n*_]. Unlike MI, the definition of directed information limits the support of *X* in the right-side term of equation 11 to the current time *i* instead of extending the support over the future time points of *X*. This term is called the ‘causal conditional entropy’ in analogy with conditional entropy. Lastly, in the absence of a causal influence of *X* on *Y*, the causal conditional entropy reduces to an entropy estimation, and therefore, *I*(*X*^*n*^ → *Y* ^*n*^) = 0.

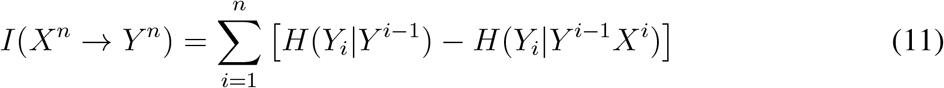

Causal conditioning might involve more than one process. For instance, in neuronal networks, the information flow between two neurons could be mediated by other neurons. Therefore, the contribution from these other time series is included as ‘side information’ while deciphering direct causal influences between pairs of neurons. Therefore, the causal conditional directed information for three processes *X, Y*, and *Z* is defined as in equation (12). Here, the number of ‘side variables’ can be greater than one; however, we incorporate only one other time series for our studies.

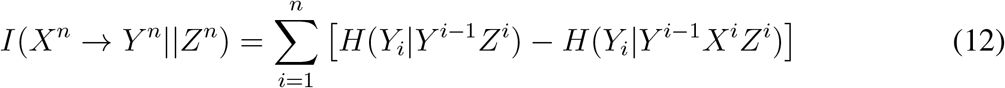

Computation of directed information requires apriori knowledge about the joint probability distribution function of the random variables. However, such information is unknown in most neuroscience scenarios of ensemble-recorded electrophysiological signals. For our studies on effective connectivity, we adopt an approach that substitutes any estimation of the probability distribution function from the data with functionals. Here, we exploit the properties of a Hilbert space approach of the entropy functional proposed in (*40*). In (*40*), the authors show that the entropy functional fulfills similar axiomatic properties of the Renyi’s *α*-order entropy (*α* > 0) (*90*), which is a generalization over the Shannon entropy. Entropy estimation using the functional entails three steps: first, the sample variable *X* = {*x*_1_, *x*_2_, …, *x*_*T*_ }, is projected onto a reproducing kernel Hilbert space (RKHS) using a positive definite kernel, *κ* : *𝒳* × *𝒳* ↦ ℝ. Next, we construct a normalized Gram matrix, *A*, on the projected data where 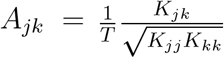, and finally, the estimator defines entropy over the eigenspectrum of the normalized Gram matrix, *A* as in equation (13), where *λ*_*j*_(*A*) denotes the *j*^*th*^ eigenvalue of *A*.

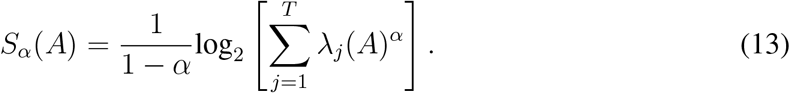

In essence, the kernel-induced mapping of the sample data into the RKHS provides a means for computing the high-order statistics of the data, wherein the eigenspectrum of the estimated Gram matrix quantifies the variations in these higher-order moments. For evaluations of joint entropy, an extension of the matrix-based entropy measure uses Hadamard products to convey the joint representation of two random variables. In this extension, joint entropy between two random variables, *X* and *Y*, is defined as in equation 14, where *A*_*x*_ and *A*_*y*_ are normalized Gram matrices evaluated on *X* and *Y*, respectively using positive definite kernels, *κ*_1_ and *κ*_2_.

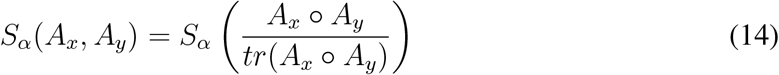

Finally, using the above estimator of entropy, directed information can be rewritten as in equations (15-16), where the latter is obtained by expanding the conditional entropy terms.

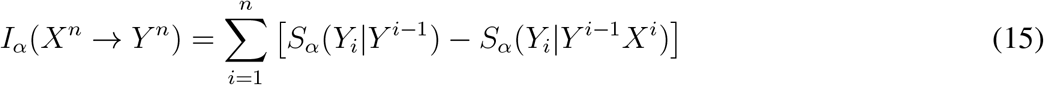

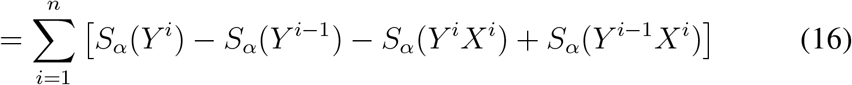

While the above approach provides an elegant yet computationally efficient formulation of directed information, we are still faced with the challenge of defining a positive-definite kernel for the point processes of spike trains. Owing to the popularity of neural spike trains, several methods have been proposed to introduce basic structure to the point process space; we focus on methods that utilize kernel constructs. Here, we briefly summarize the approach. Firstly, each point process is brought into the continuous *L*_2_ space as intensity functions via convolutional operators. Next, using an RKHS inducing kernel, *κ* : *𝒳* × *𝒳* ↦ ℝ and evoking the kernel trick, we evaluate the normalized Gram matrix, *A*, as the pairwise inner product of the intensity functions in the Hilbert space.i.e., ⟨*x*_1_, *x*_2_⟩_ℌ_ = *κ*(*x*_1_, *x*_2_). In this way, we obtain *A* as a symmetric positive semidefinite matrix. A complete schematic demonstrating the evaluation of the joint entropy term, *S*_*α*_ (*Y*^*i*^, *X*^*i*^), is summarized in Figure 1**B**, where the time-shifted matrices, *T*_*x*_ and *T*_*y*_, capture observations from each intensity function for DI estimation.

For projections into the RKHS, we employ the non-linear Schoenberg kernels (*κ*) (with a rectangular window for intensity function) as the most appropriate choice. Firstly, Schoenberg kernels are positive definite kernels - a necessary property for formulating the matrix-based entropy. Second, derived from radial basis functions, these kernels are universal and can provably approximate arbitrary non-linear functions from the point process to the real space. Third, the locality of a rectangular intensity function limits the causal measure to events only within the window reflective of the system’s memory under study. Finally, in a rigorous simulation study conducted by us (*91*), these kernels outperformed other positive definite kernels in identifying the correct causal direction. The complete formulation of the kernel for two intensity functions, *x* and *y*, is shown in equations (17 - 18). Here, 𝕀 is the indicator function, *w* is the width of the rectangular window, *σ*_*κ*_ is the kernel size, and 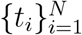 are the event timings.

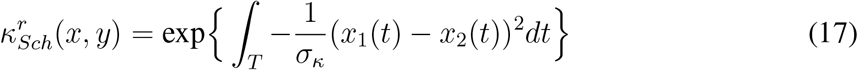

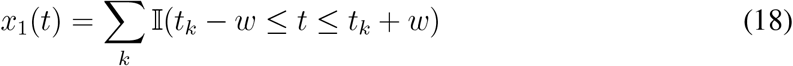

Overall, we utilize the Hilbert space as an extended functional representation for point process analysis enabling the estimation of directed information for point process systems. Finally, we note a critical consequence of the estimation of directed information. Although theoretically, the directed information between independent time series is equal to zero, in practice, empirical values are small and non-negative. We address this issue via appropriate significance testing for the signals under analysis. The MATLAB implementation of our DI model can be found at https://github.com/shailajaAkella/Directed-Information.

#### Parameter selection

Our formulation of directed information depends on four hyperparameters: 1) the kernel size (*σ*_*κ*_), 2) the width of the rectangular window (*w*), 3) the order *α*, and 4) memory of the DI measure (*n*). The kernel size dictates the geometrical extent of the inner product in the RKHS. In all our analyses, the selection of kernel size follows Scott’s rule of thumb for density estimation (*92*). It is important to note that for projections into the same RKHS, the kernel size must be the same for all entropy evaluations. In our studies, this was chosen as the kernel size of the input causal variable. Next, for each point process of beta, gamma, and spiking activity, the width of the rectangular intensity operator was fixed as 250 ms, 120 ms, and 120 ms, respectively, to approximately cover the duration of the oscillatory burst events. Third, the choice of *α* is associated with the task under study. For emphasis on rare events (i.e., in the tails of the distribution), *α* must be chosen close to 1 (*40*), while higher values of *α* (> 2) characterize modal behavior. In our analysis of effective connectivity, we are interested in sparse transient synchronizations between multiscale activity. Therefore, we set the order *α* to 1.01. Lastly, to capture signal transmissions occurring at brief timescales, the DI measure’s memory, *n*, was chosen to be 20 ms. Since each row in the time-shifted matrix (Figure 1**B**) maximally depends on time delays captured by the rectangular window used for intensity evaluation and the DI memory, the DI estimations culminate in total time dependency of *w* + *n*. Therefore, our metric evaluates time dependencies of 270 ms for beta and 140 ms for gamma and spiking activities.

### Constructing Causal Graphs

Connectivity between brain structures or ‘nodes’ can be best visualized via a graphical representation. In our connectivity studies, a ‘node’ corresponds to an individual neuron (spikes) or neuronal population (fields), the edges between the nodes refer to the connectivity patterns formed by causal influences, and the directionality of the causal influence is represented as a directed edge between two nodes. To this end, we have constructed a causal measure that quantifies directed information flow between the cortical nodes whose properties are characterized via a point process. Since, in practice, all estimated DI measures (between causally and non-causally connected nodes) are greater than zero, we have quantified a fully connected directed graph between all observed nodes.

The next step is to identify the edges that significantly influence causation. However, this is challenging because significance testing requires knowledge of the DI measure’s probability density function (pdf) under the null hypothesis (non-causality). It is difficult to pin down the null hypothesis distribution due to the statistical variability of neural activity. Next, we must account for recurrent connections within single-neuron circuits. At the same time, we also need to consider that spike-field interactions have a unidirectional causal effect at a particular time due to the physics of wave propagation in the brain media. To handle these differences and difficulties, we designed separate significance testing pipelines for each interaction type.

#### Spike-Spike interactions

To test the significance of the DI measures for spike-spike interactions, we implemented a random permutation procedure to build a baseline null hypothesis distribution. Next, we compared the evaluated DI measure on the actual data against this baseline distribution to assess the significance level of the DI measure. Specifically, we create a surrogate system by shuffling the spike times of the causal variable. Therefore, the shuffled point process maintains the spike counts in each realization, but any consistent structure in the joint probability distribution between the two-time series is lost. Performing such random shuffling with many different permutations results in a DI distribution corresponding to the null hypothesis of non-causality. We reject the null hypothesis if the DI values estimated on the original time series are significantly greater than the baseline threshold at a level of *α*_*s*_ = 0.05.

However, we have yet to interpret the connectivity patterns meaningfully as neuronal nodes are recurrently connected, creating copious amounts of indirect influences. Indirect influences can be teased apart from direct influences by conditioning on ‘side processes’ whose knowledge renders the involved processes statistically independent. Two types of indirect influences considered in this study are the ‘proxy’ and ‘cascading’ influences (Figure 2**A** right). In a proxy influence pattern, process *Y*_1_ influences process *Y*_2_ which in turn drives the process *Y*_3_ (Figure 2**A** top-right). In such a case, the DI measure will likely identify an indirect influence of process *Y*_1_ on *Y*_3_. However, the two processes can be rendered statistically independent given causal knowledge of *Y*_2_ → *Y*_3_. Similarly, in a cascading topology where two processes, *Y*_1_ and *Y*_3_, are commonly driven by a third process, *Y*_2_, the indirect connectivity between the two could be accounted for by their dependence on *Y*_2_ (Figure 2**A** bottom-right).

Causal conditioning of DI on ‘side processes’ is performed as shown in equation (12) such that *X* directly, causally influences *Y*, iff *I*(*X*^*n*^ → *Y* ^*n*^||*Z*^*n*^) > 0. Nevertheless, the problem of significance testing is also critical to this analysis as the null hypothesis distribution of the conditional DI measure is also changing, and their estimates between independent processes never typically equate to zero. Therefore, to classify the direct connectivities in the network, we evaluate the decrease in estimated influence between two nodes after introducing a side variable. Firstly, potential indirect influence patterns are identified as three nodes that create cascading and proxy-type dependencies in the directed graph. Across all combinations of existing dependencies between the three nodes, we evaluate the percentage change in the DI measure upon conditioning on the third node. Across edges terminating on the same node, any edge that registers a significantly larger change in the DI measure rejects the null hypothesis that all connecting edges to the terminating node directly influence it. The edge with the largest change is classified as indirect and removed from the network.

#### Spike-field interactions

Unlike spike-spike interactions, we assume a unidirectional causal influence between spikes and fields. Under this assumption, it is sufficient to compare the DI estimates of the opposing influences, i.e., between DI(spikes → fields) and DI(fields → spikes), to identify the dominant connection between the signal modalities. Moreover, we account for the effect of large sample size on significance testing by using both substantive and statistical significance measures to identify causal influences (*93*). First, we test the hypothesis that the DI measures from the opposing connections are drawn from populations with different means against the null hypothesis that the population means are the same. If the test rejects the null hypothesis at the 1% significance level, we check for substantive significance by evaluating the effect size (*η*) between the DI measures of the two connectivity groups. To estimate the effect size, we use the Cohen’s - d coefficient (equation 20) where ∆*DI* is the difference between the group means, and *s*_1_ and *s*_2_ are the groups’ standard deviations. Significant causal influences are determined as those that register effect sizes greater than 0.2, qualifying those connections that report medium to large differences between the two groups (*94*).

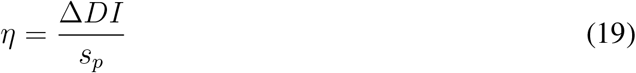

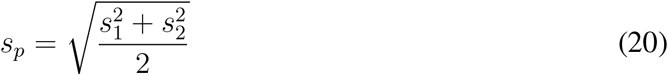

(19)

### Validation

We compared the performance of our DI model with Granger causality (GC) (*95*) on a six-neuron network model comprising indirect influences, explicit driver neurons, single input neurons, different synaptic transmissions, and simple excitatory connections. To test these models, we used the Izhikevich neuron model to simulate neuronal activity for all neurons. The Izhikevich model (*96*) is a mathematical model that can reproduce the general behavior of the more biophysically accurate Hodgkin-Huxley-type models using a computationally simpler and more efficient formulation. The model is defined by a system of two differential equations and several parameters that dictate spiking properties. The Izhikevich neuron model is governed by the following equations (equations 8.5 and 8.6 in (*96*)),

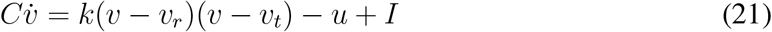

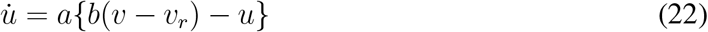

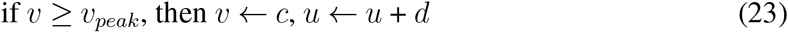

Here, *v* is the neuron’s membrane potential, and *u* is the membrane recovery variable. The latter is associated with activation and inactivation of *K*^+^ and *Na*^+^ ionic currents, respectively, that provide negative feedback to the membrane potential. Description of the hyperparameters and their corresponding values for a regular-spiking neuron are listed in Table S1.

Each neuron, 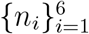, could be simulated using three types of input current, *I*_*in*_: 1) constant input, 2) uniform input, a uniformly distributed value between (0 − 150) pA, or 3) bursty input, two 150 ms long pulses of 300 pA, each supplied at a random time, interleaved with a uniformly distributed current between (0−150) pA (Figure S4**A, B**). The latter input was supplied to create a system of bursty neurons where only the presynaptic neuron was stimulated in this manner. The postsynaptic neuron, however, received an input of the former type. Specifically, uniform input current was supplied to neurons *n*_1_, *n*_2_, *n*_3_ and *n*_6_. Whereas *n*_5_ received a constant input current of 150 pA, *n*_4_ was simulated to mimic an interneuron such that it received a bursty input current. The magnitude of excitation current, *I*_*e*_ (or inhibition current *I*_*i*_) from an excitatory neuron (or inhibitory neuron) in a connection was kept constant throughout each simulation. These values were set at *I*_*e*_ = 350 pA and *I*_*i*_ = 180 pA. The total current, *I*, supplied to each neuron comprised of *I*_*in*_ and *I*_*i/e*_. We generated membrane noise as Gaussian noise and combined it with the neuron’s membrane potential, *v*, at each time point.

Network connections were made to create a total of 5 direct excitatory driver connections connections: *n*_1_ → *n*_6_, *n*_2_ → *n*_1_, *n*_2_ → *n*_3_, *n*_5_ → *n*_4_, and *n*_5_ → *n*_6_. The network also included an inhibitory connection between *n*_4_ → *n*_2_. This network topology, thus, created several firstorder indirect connections deriving from ‘proxy’ (A → B, B → C) and ‘cascading’ (A → B, A → C) influences in the network. Indirect connections that derived from a ‘proxy’ topology included: *n*_2_ → *n*_6_, *n*_4_ → *n*_1_, *n*_5_ → *n*_2_, and *n*_4_ → *n*_3_. Whereas, indirect connections that derived from a ‘cascading’ topology included: *n*_1_ ↔ *n*_3_, *n*_1_ ↔ *n*_5_, and *n*_6_ ↔ *n*_4_ (Figure S4**B**). In the network, synaptic delay between two connected neurons was maintained at 5 ms. Accordingly, the memory for all DI evaluations was set at 20 ms, same as the order of the filters in the GC model. Finally, we simulated 100 trials with a time-resolution of 1 ms such that each trial lasted for 1s.

To identify significant connections using the GC model, we followed the methods shown in (*95*) for evaluating the likelihood ratio as well significance testing. Specifically, each edge was placed under a *χ*^2^ test as well as a false discovery ratio test to correct for multiple comparisons. Both the DI and GC models were able to identify all direct causal patterns where the adjacency matrix comprising all identified direct connections are presented in Figure S4**C, D**, respectively. While the DI model achieved a connectivity detection accuracy of 96.67%, the detection accuracy of the GC model was 80%. The overall neuronal network includes many potentially indirect influences. Yet, the DI model only identifies three indirect edges, demonstrating high robustness to spurious causal influences (Figure S4**C**, middle). Post-significance testing for indirect connections, DI correctly identified two of the three as indirect (Figure S4**C**, right), a false positive rate of 3%. Inquiries into the remaining edge from *n*_3_ to *n*_6_ revealed that this connection was never checked for indirect influences. The model’s current implementation only checks for indirect causal connections between nodes involved in indirect dependencies of length 3. In the present example, neither {*n*_1_, *n*_3_, *n*_6_} nor {*n*_5_, *n*_3_, *n*_6_} form an undirected cycle among themselves in the estimated connectivity map (Figure S4**B**). Therefore, these interactions were not considered indirect by the model. It is important to note that our current implementation of the causal graph construction is easily extendable to include verifications on more dependencies. On the other hand, the GC model detected many more indirect connections, leading to a false positive rate of 20% (Figure**D**).

## Acknowledgements

The authors declare that they have no affiliations with or involvement in any organization or entity with any commercial or financial interest that could be construed as a potential conflict of interest. This work was supported by the NSF 1631759 (J.C.P), Office of Naval Research N00014-22-1-2453 (E.K.M), The Picower Institute for Learning and Memory (E.K.M), The JPB Foundation (E.K.M.) and National Institute of Mental Health NIMH 1R01MH131715-01 (E.K.M).

## Ethics Statement

All procedures involving animals were reviewed and approved by the Massachusetts Institute of Technology (MIT)’s Committee on Animal Care and were conducted in accordance with the guidelines of the National Institute of Health and MIT’s Department of Comparative Medicine.

## Supplementary materials

Figs. S1 to S11

Tables S1

## References

1. S. van Pelt, et al., Social cognitive and affective neuroscience 11, 973 (2016).

2. A. M. Bastos, M. Lundqvist, A. S. Waite, N. Kopell, E. K. Miller, Proceedings of the National Academy of Sciences 117, 31459 (2020).

3. G. G. Gregoriou, S. J. Gotts, H. Zhou, R. Desimone, science 324, 1207 (2009).

4. A. M. Bastos, et al., Neuron 85, 390 (2015).

5. C. G. Richter, R. Coppola, S. L. Bressler, Scientific reports 8, 1 (2018).

6. D. Ferro, J. van Kempen, M. Boyd, S. Panzeri, A. Thiele, Proceedings of the National Academy of Sciences 118, e2022097118 (2021).

7. T. Popov, O. Jensen, J.-M. Schoffelen, NeuroImage 178, 277 (2018).

8. V. Dimakopoulos, P. Mégevand, L. H. Stieglitz, L. Imbach, J. Sarnthein, Elife 11, e78677 (2022).

9. K. Tanaka, Journal of neurophysiology 49, 1303 (1983).

10. J.-M. Alonso, W. M. Usrey, R. C. Reid, Nature 383, 815 (1996).

11. Z. Wang, et al., Neuron 78, 1116 (2013).

12. J. H. Siegle, et al., Nature 592, 86 (2021).

13. X. Jia, et al., Neuron 110, 1585 (2022).

14. J. D. Semedo, et al., Nature communications 13, 1 (2022).

15. I. H. Stevenson, J. M. Rebesco, L. E. Miller, K. P. Körding, Current opinion in neurobiology 18, 582 (2008).

16. M. R. Cohen, A. Kohn, Nature neuroscience 14, 811 (2011).

17. C. Clopath, L. Büsing, E. Vasilaki, W. Gerstner, Nature neuroscience 13, 344 (2010).

18. W. J. Freeman, Journal of Physiology-Paris 94, 303 (2000).

19. G. Buzsaki, A. Draguhn, science 304, 1926 (2004).

20. S. Ray, Current opinion in neurobiology 31, 111 (2015).

21. D. J. Felleman, D. C. Van Essen, Cerebral cortex (New York, NY: 1991) 1, 1 (1991).

22. A. M. Thomson, D. C. West, Y. Wang, A. P. Bannister, Cerebral cortex 12, 936 (2002).

23. J. Maunsell, D. C. van Essen, Journal of Neuroscience 3, 2563 (1983).

24. R. Desimone, T. D. Albright, C. G. Gross, C. Bruce, Journal of Neuroscience 4, 2051 (1984).

25. C. E. Connor, S. L. Brincat, A. Pasupathy, Current opinion in neurobiology 17, 140 (2007).

26. C. A. Atencio, T. O. Sharpee, C. E. Schreiner, Proceedings of the National Academy of Sciences 106, 21894 (2009).

27. C. A. Atencio, C. E. Schreiner, PLoS One 5, e9521 (2010).

28. Y. Iwamura, Current opinion in neurobiology 8, 522 (1998).

29. S. Lefort, C. Tomm, J.-C. F. Sarria, C. C. Petersen, Neuron 61, 301 (2009).

30. N. T. Markov, H. Kennedy, Current opinion in neurobiology 23, 187 (2013).

31. X.-J. Wang, Physiological reviews 90, 1195 (2010).

32. R. J. Douglas, K. Martin, The Journal of physiology 440, 735 (1991).

33. K. Friston, Neural Networks 16, 1325 (2003).

34. K. Friston, PLoS computational biology 4, e1000211 (2008).

35. K. Friston, Nature reviews neuroscience 11, 127 (2010).

36. E. A. Buffalo, P. Fries, R. Landman, T. J. Buschman, R. Desimone, Proceedings of the National Academy of Sciences 108, 11262 (2011).

37. S. Akella, A. Mohebi, J. C. Principe, K. Oweiss, Journal of Neural Engineering 18, 026016 (2021).

38. A. M. Bastos, R. Loonis, S. Kornblith, M. Lundqvist, E. K. Miller, Proceedings of the National Academy of Sciences 115, 1117 (2018).

39. M. Lundqvist, A. M. Bastos, E. K. Miller, Journal of cognitive neuroscience 32, 2024 (2020).

40. L. G. S. Giraldo, M. Rao, J. C. Principe, IEEE Transactions on Information Theory 61, 535 (2014).

41. A. Maier, G. K. Adams, C. Aura, D. A. Leopold, Frontiers in systems neuroscience 4, 31 (2010).

42. A. K. Roopun, et al., Frontiers in cellular neuroscience p. 1 (2008).

43. T. J. Buschman, E. K. Miller, science 315, 1860 (2007).

44. X. Gong, W. Li, H. Liang, Journal of neurophysiology 122, 809 (2019).

45. P. L. Nunez, R. Srinivasan, et al., Electric fields of the brain: the neurophysics of EEG (Oxford University Press, USA, 2006).

46. U. Mitzdorf, Physiological reviews 65, 37 (1985).

47. N. K. Logothetis, B. A. Wandell, Annu. Rev. Physiol. 66, 735 (2004).

48. R. Elul, International review of neurobiology 15, 227 (1972).

49. G. Buzsáki, Nature neuroscience 7, 446 (2004).

50. O. Jensen, Neuroscience 139, 237 (2006).

51. R. T. Canolty, et al., Proceedings of the National Academy of Sciences 107, 17356 (2010).

52. E. Salinas, T. J. Sejnowski, Nature reviews neuroscience 2, 539 (2001).

53. P. Fries, Trends in cognitive sciences 9, 474 (2005).

54. M. J. Rasch, A. Gretton, Y. Murayama, W. Maass, N. K. Logothetis, Journal of neurophysiology 99, 1461 (2008).

55. O. Herreras, Frontiers in neural circuits 10, 101 (2016).

56. M. T. Schmolesky, et al., Journal of neurophysiology 79, 3272 (1998).

57. D. Jokisch, O. Jensen, Journal of Neuroscience 27, 3244 (2007).

58. M. Lundqvist, et al., Neuron 90, 152 (2016).

59. P. Fries, J. H. Reynolds, A. E. Rorie, R. Desimone, Science 291, 1560 (2001).

60. M. Bauer, R. Oostenveld, M. Peeters, P. Fries, Journal of Neuroscience 26, 490 (2006).

61. E. K. Miller, M. Lundqvist, A. M. Bastos, Neuron 100, 463 (2018).

62. H. Choi, A. Pasupathy, E. Shea-Brown, Neural computation 30, 1209 (2018).

63. T. Grent-’t Jong, M. G. Woldorff, PLoS biology 5, e12 (2007).

64. J. A. Harris, et al., Nature 575, 195 (2019).

65. A. M. Bastos, et al., Neuron 76, 695 (2012).

66. E. M. Callaway, Annual review of neuroscience 21, 47 (1998).

67. D. Xing, C.-I. Yeh, S. Burns, R. M. Shapley, Proceedings of the National Academy of Sciences 109, 13871 (2012).

68. J. Vezoli, et al., Neuron 109, 3862 (2021).

69. S. Paneri, G. G. Gregoriou, Frontiers in neuroscience 11, 545 (2017).

70. R. P. Rao, D. H. Ballard, Nature neuroscience 2, 79 (1999).

71. M. W. Spratling, Frontiers in computational neuroscience p. 4 (2008).

72. H. R. Brown, K. J. Friston, Frontiers in human neuroscience 7, 784 (2013).

73. L. H. Arnal, A.-L. Giraud, Trends in cognitive sciences 16, 390 (2012).

74. L. Zhang, et al., Hippocampus 22, 1781 (2012).

75. D. Kang, M. Ding, I. Topchiy, B. Kocsis, Frontiers in neuroanatomy 11, 120 (2017).

76. H.-L. Hsieh, Y. T. Wong, B. Pesaran, M. M. Shanechi, Journal of neural engineering 16, 016018 (2018).

77. C. Wang, B. Pesaran, M. M. Shanechi, Journal of neural engineering 19, 026001 (2022).

78. D. Gabor, Journal of the Institution of Electrical Engineers-part III: radio and communication engineering 93, 429 (1946).

79. S. M. Sherman, R. Guillery, Journal of neurophysiology 106, 1068 (2011).

80. Y. B. Saalmann, M. A. Pinsk, L. Wang, X. Li, S. Kastner, science 337, 753 (2012).

81. W. J. Freeman, et al., Mass action in the nervous system, vol. 2004 (Citeseer, 1975).

82. E. Niedermeyer, F. L. da Silva, Electroencephalography: basic principles, clinical applications, and related fields (Lippincott Williams & Wilkins, 2005).

83. W. B. Davenport, W. L. Root, et al., An introduction to the theory of random signals and noise, vol. 159 (McGraw-Hill New York, 1958).

84. S. Akella, J. C. Principe, 2019 41st Annual International Conference of the IEEE Engineering in Medicine and Biology Society (EMBC) (IEEE, 2019), pp. 5790–5793.

85. S. Akella, et al., 2021 10th International IEEE/EMBS Conference on Neural Engineering (NER) (IEEE, 2021), pp. 420–425.

86. B. W. Silverman, Density estimation for statistics and data analysis (Routledge, 2018).

87. M. Little, P. Mcsharry, S. Roberts, D. Costello, I. Moroz, Nature Precedings pp. 1–1 (2007).

88. M. J. Prerau, R. E. Brown, M. T. Bianchi, J. M. Ellenbogen, P. L. Purdon, Physiology 32, 60 (2017).

89. P.-O. Amblard, O. J. Michel, Entropy 15, 113 (2012).

90. A. Rényi, et al., Proceedings of the fourth Berkeley symposium on mathematical statistics and probability (Berkeley, lCalifornia, USA, 1961), vol. 1.

91. S. Akella, J. C. Principe, Neuromodulatory pattern analysis for local field potentials, Ph.D. thesis, University of Florida (2021).

92. D. W. Scott, S. R. Sain, Handbook of statistics 24, 229 (2005).

93. G. M. Sullivan, R. Feinn, Journal of graduate medical education 4, 279 (2012).

94. J. Cohen, Current directions in psychological science 1, 98 (1992).

95. S. Kim, D. Putrino, S. Ghosh, E. N. Brown, PLoS computational biology 7, e1001110 (2011).

96. E. M. Izhikevich, Dynamical systems in neuroscience (MIT press, 2007).

