## Supplementary figures for "Measurable fields-to-spike causality and its dependence on cortical layer and area"

Shailaja Akella *et al.*

#### Supplementary materials

This PDF file includes:

1. Figures S1 - S11

2. Table S1

### Figures

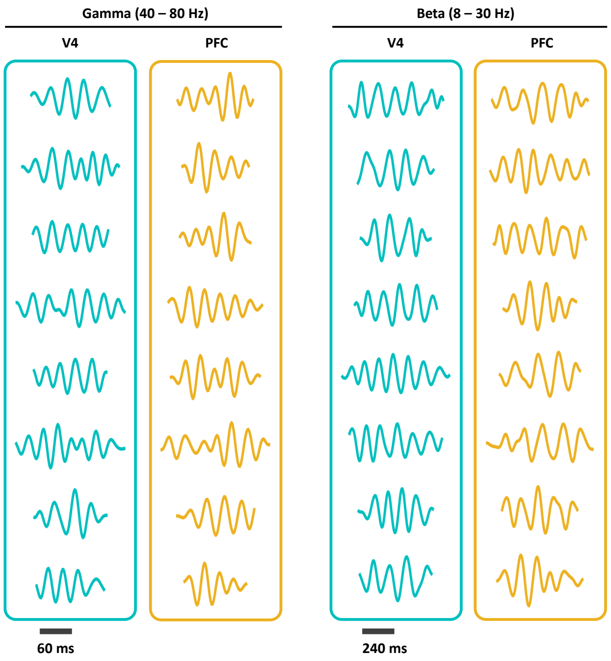


Figure S1: Example waveforms from the dictionary constructed using the generative model in the gamma and beta frequency ranges in the areas, V4 and PFC.

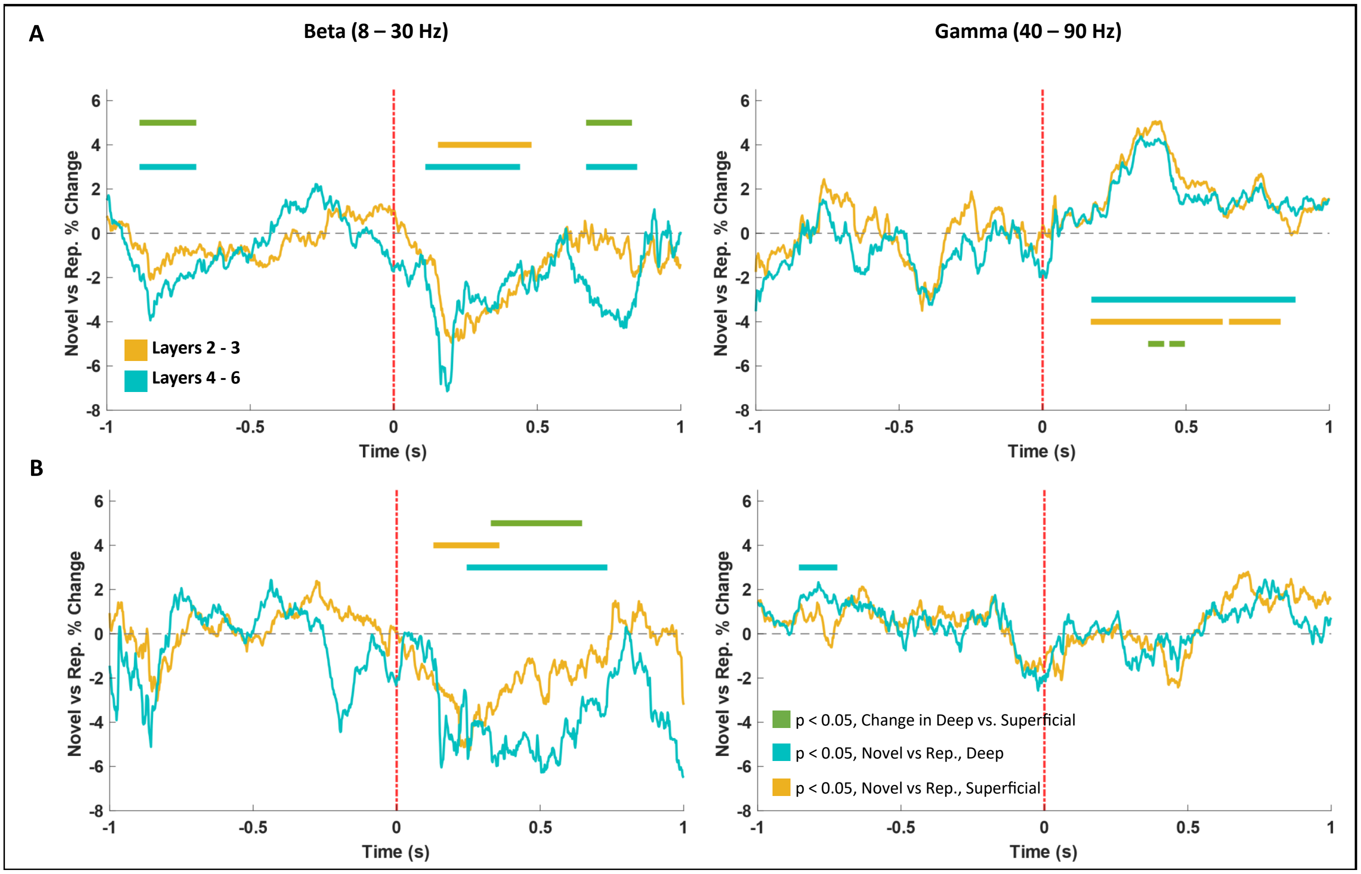


Figure S2: Change in instantaneous beta (column 1) and gamma (column 2) bursting power between randomized and repetitive trial blocks in (**A**) V4 and (**B**) PFC. Signals were analyzed during fixation, [-1, 0]s and stimulus processing [0, 1]s. Data are mean ± sem.

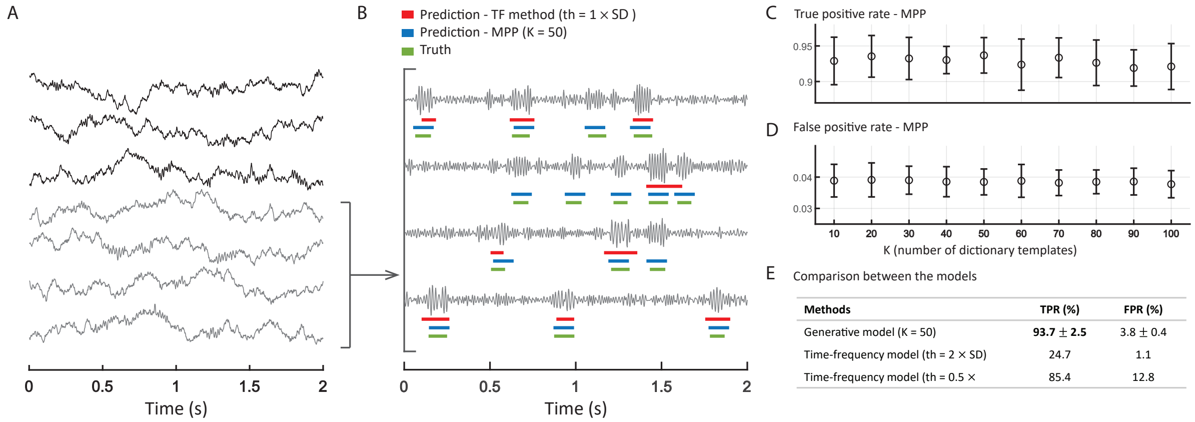


Figure S3: Validation of the generative model for oscillatory bursts. **A**, Example LFP data across seven trials. **B**, Filtered LFP between 40-80 Hz demarcating the time points of true bursts (green), and bursts obtained via the generative model (MPP, blue) and amplitude thresholding (TF, red). **C, D**, True positive rate (TPR) and false positive rate (FPR) for the generative model. **E**, Comparison of model performance (TPR, FPR) between the best generative and thresholding models.

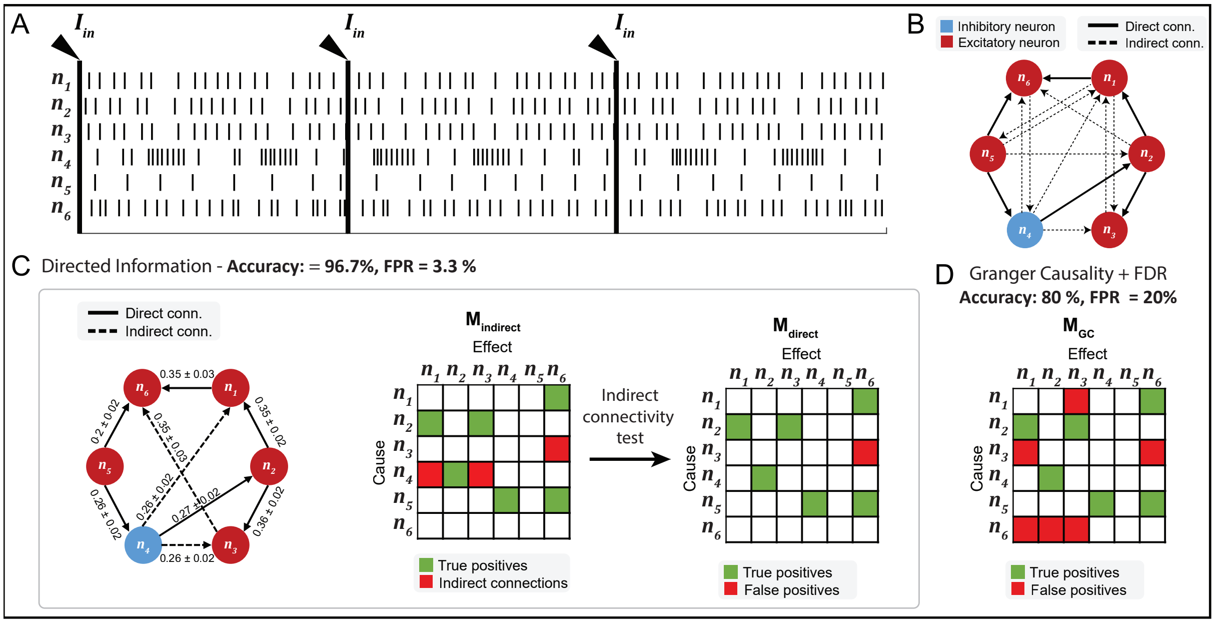


Figure S4: Identification of direct causal networks in a multi-neuron system using DI and Granger causality (GC). **A**, Example data presenting 3 trials in each neuron. **B**, Network connectivity pattern. **C**, Results from DI model. Left, DI estimates for all directions identified by the DI model. Data are mean ± std. Solid lines correspond to true direct causal directions, while dashed lines are indirect causal influences. Middle, Estimated adjacency matrix including all causal influences. Right, Estimated adjacency matrix including only direct causal influences. **D**, Final adjacency matrix estimated using the GC model.

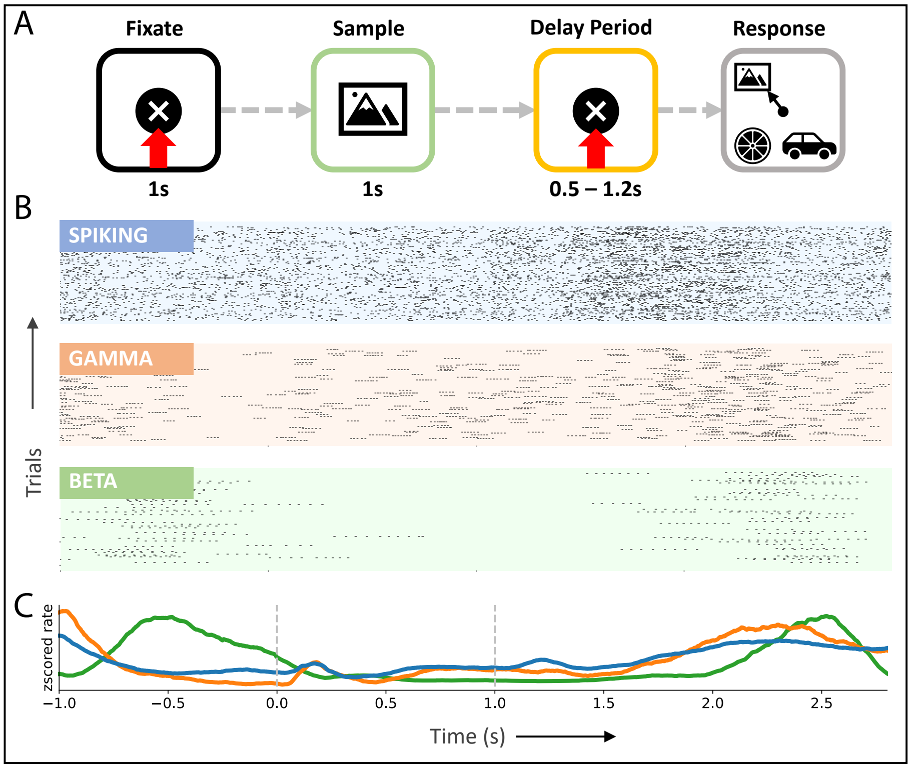


Figure S5: **A** Experimental setup. Following a 1s-fixation period, the subject is shown a sample stimulus for 1s. At the end of a delay period (fixed or variable), the subject saccades to the sampled stimulus that reappears at one of four randomized locations along with distractor images. **B**, Exemplar spiking activity and LFPs from layer 2/3 of PFC visualized in the same space of point processes, subject 2. **C**, Multi-scale average rate plots obtained via kernel smoothing methods, subject 2. All activity is aligned with the trial intervals in **C**.

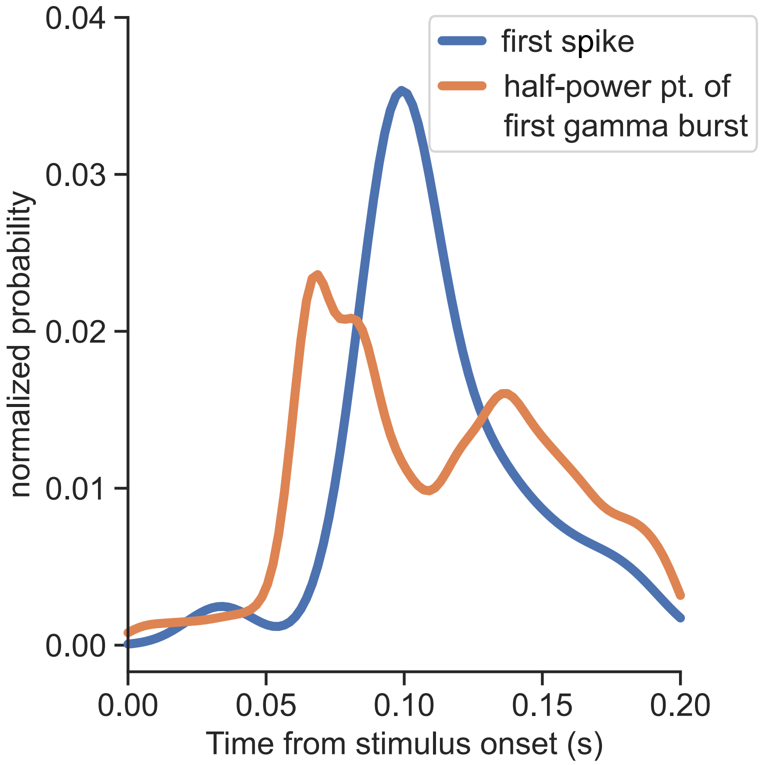


Figure S6: Distribution of time to first spike and half-power point of the first gamma burst in response to sample stimulus across all units and layers in V4.

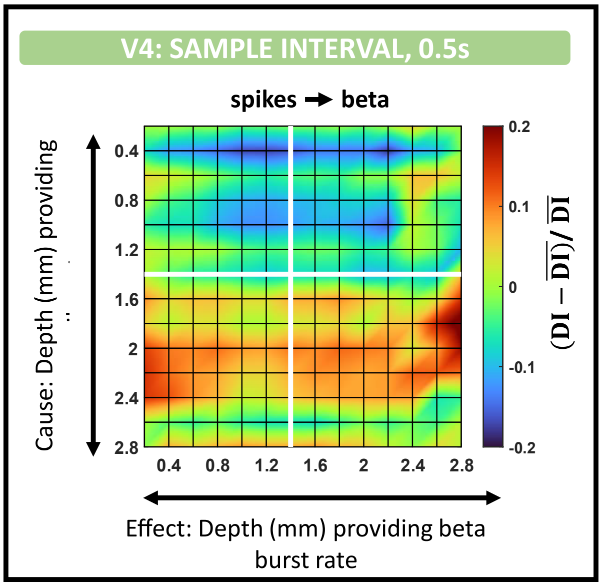


Figure S7: Spike → beta layer-wise connectivity plot in V4 during early sample interval.

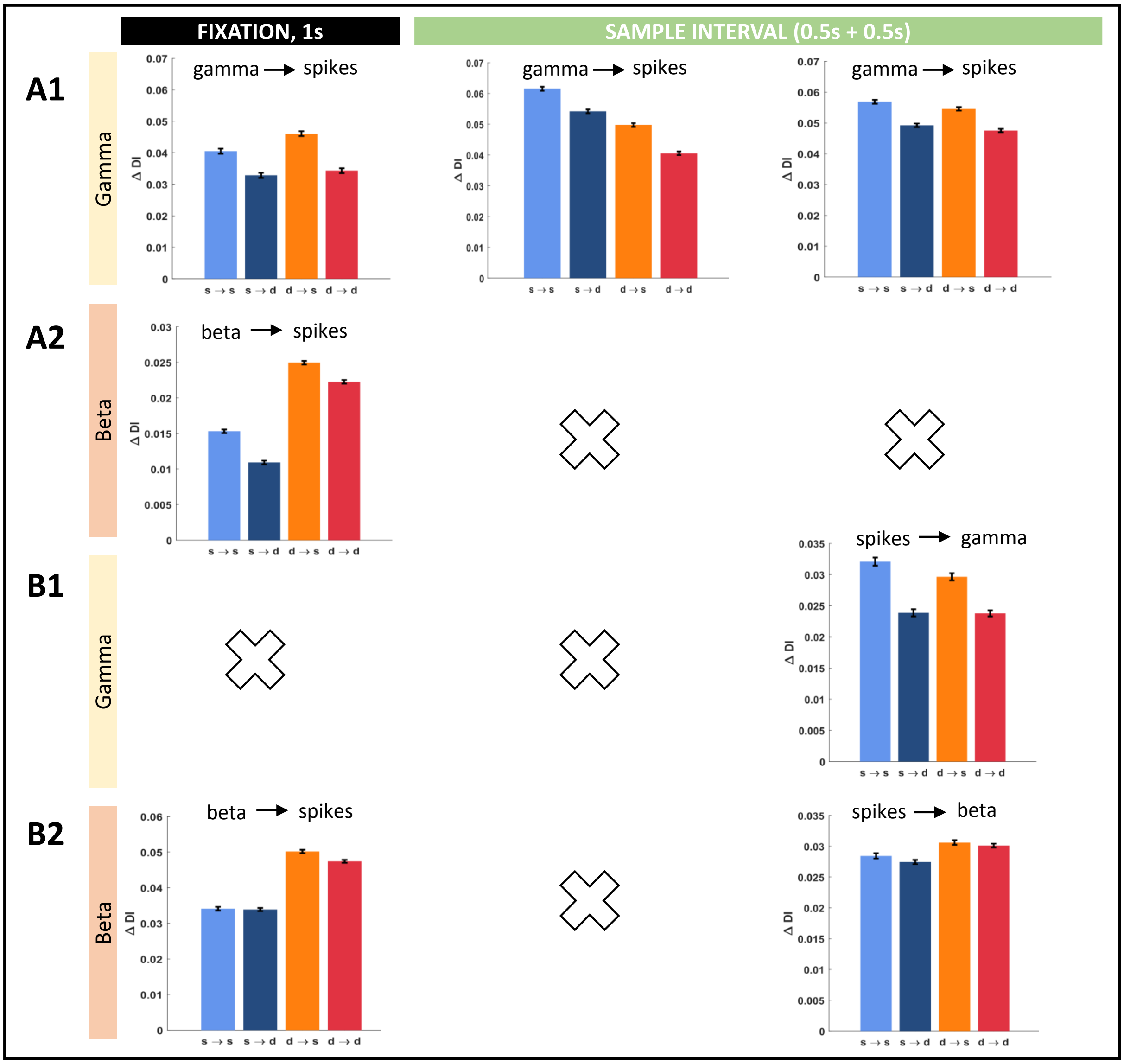


Figure S8: Summary of intralaminar spike-field connectivity patterns in **A** V4 and **B** PFC over fixation and stimulus presentation. Data are mean ± sem. ΔDI is the difference in DI between the counter-influences.

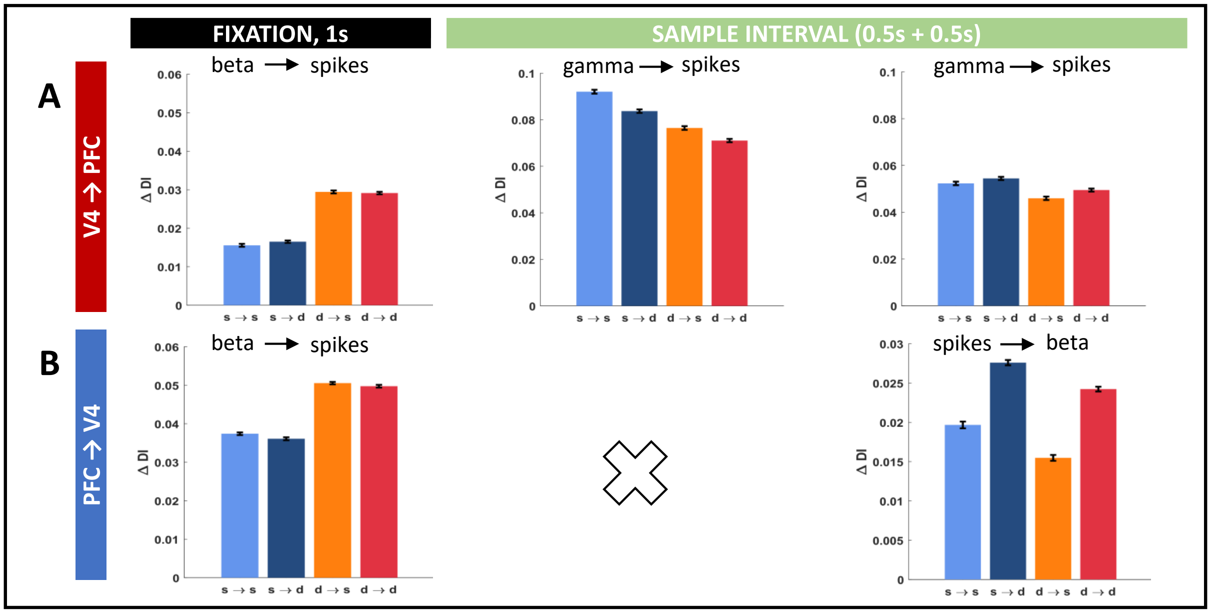


Figure S9: Summary of interlaminar spike-field connectivity patterns in the **A** feedforward (V4 $\rightarrow$ PFC) and **B** feedback (PFC $\rightarrow$ V4) directions during fixation and stimulus interval. Data are mean $\pm$ sem. $\Delta$DI is the difference in DI between the counter-influences.

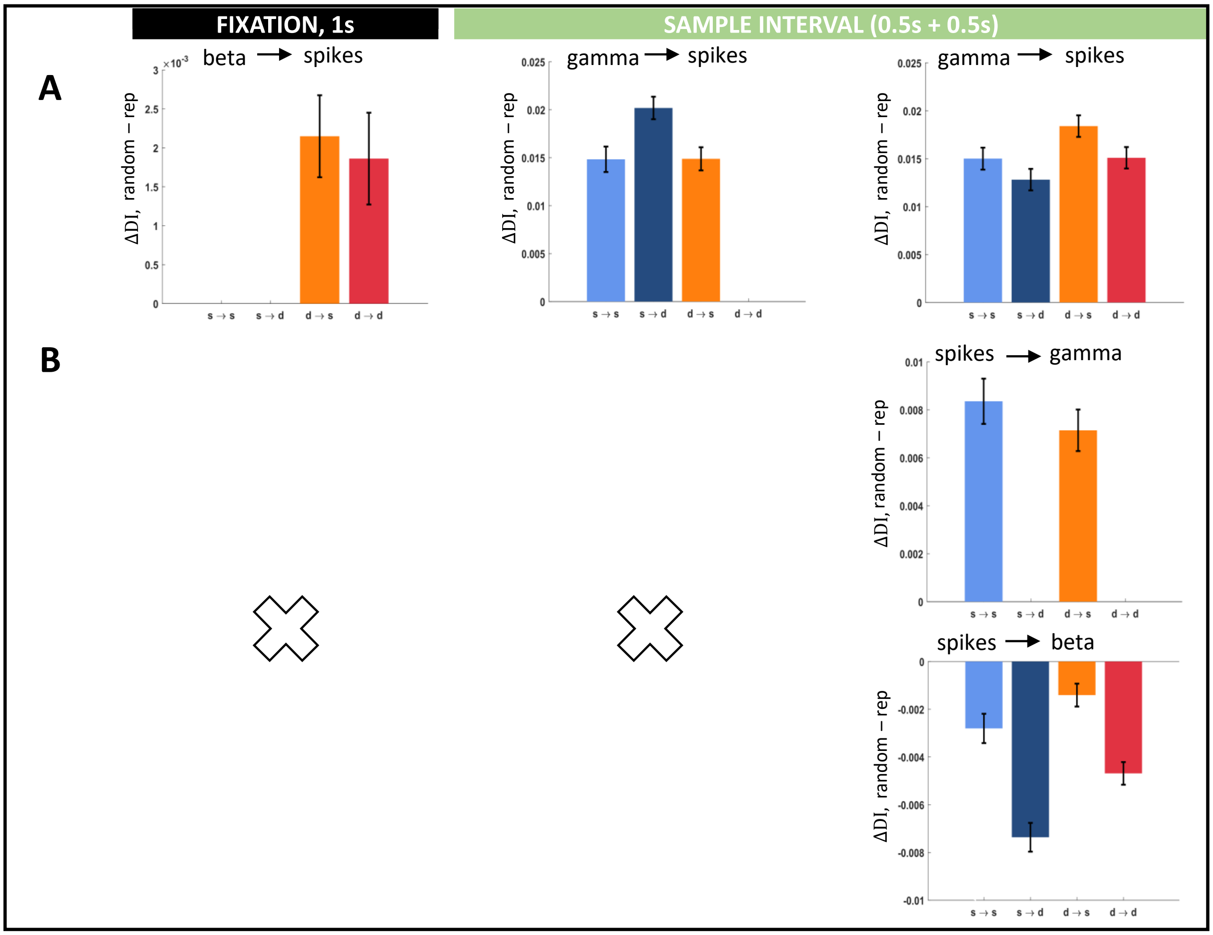


Figure S10: Effect of stimulus predictability on intralaminar spike-field connectivity patterns in the (**A**) V4 and (**B**) PFC during fixation and stimulus presentation. Data are mean $\pm$ sem. $\Delta$DI is the difference between the DI estimates in randomized and repetitive trial blocks.

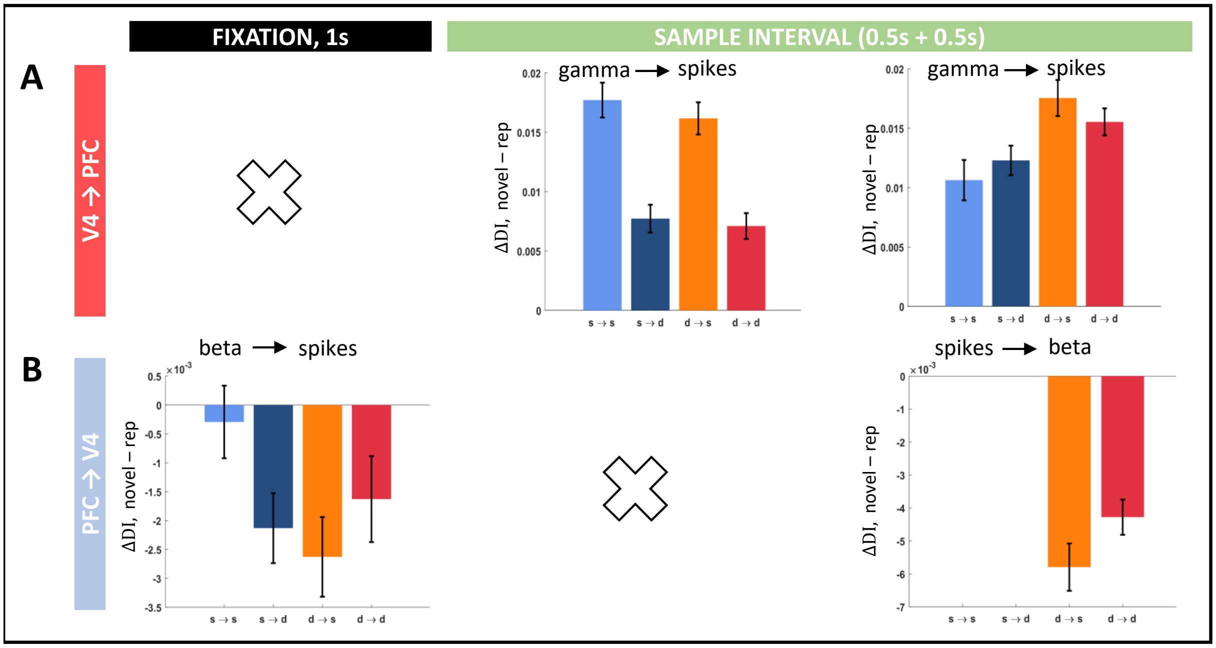


Figure S11: Effect of stimulus predictability on interlaminar spike-field connectivity patterns in the (**A**) V4 and (**B**) PFC during fixation and stimulus presentation.Data are mean ± sem. ΔDI is the difference between the DI estimates in randomized and repetitive trial blocks.

### Tables

Table S1: Izhikevich neuron model parameters for a regular spiking neuron

| **Model Parameter** | **Variable** | **Value** voltages-mV, current-pA |
|---|---|---|
| Membrane Capacitance | $C$ | 100 |
| Resting Membrane Potential | $v_r$ | -60 |
| Instantaneous Threshold Potential | $v_t$ | -40 |
| Peak Voltage | $v_{peak}$ | 35 |
| Recovery Time Constant | a | 0.03 |
| Related to neuron's rheobase and input resistance | b, k | -2, 0.7 |
| Voltage Reset Potential | c | -50 |
| Net current flow during an action potential | d | 100 |
